# Vascular development of fetal and postnatal neocortex of the pig, the European wild boar *Sus scrofa*

**DOI:** 10.1101/2024.05.17.594685

**Authors:** Eric Sobierajski, Katrin Czubay, Christa Beemelmans, Christoph Beemelmans, Martin Meschkat, Dennis Uhlenkamp, Gundela Meyer, Petra Wahle

## Abstract

The development of the brain’s vascular system is a predominantly prenatal process in mammalian species and required for neurogenesis and further brain development. Our recent work on fetal pig has revealed that many neurodevelopmental processes start well before birth and proceed rapidly reaching near-mature status already around birth. Here, we analyzed the development of neocortical vasculature from embryonic day (E) 45 onwards (gestation in pig lasts 114 days) using qualitative and quantitative image analyses and protein blots. In all cortical layers, vessel volume from total brain volume at E100 resembled that of a postnatal day (P) 30 piglet. Endothelial cells expressed the tight junction protein claudin-5 from E45 onwards. GFAP+ and AQP4+ astrocytes, PDGFRβ+ pericytes and α-SMA+ smooth muscle cells are detectable near vessels at E60 suggesting an early assembly of blood-brain barrier components. The vascular system in the visual cortex is advanced before birth with an almost mature pattern at E100. Findings were confirmed by blots which showed a steady increase of expression of tight junction and angiogenesis-related proteins (claudin-5, occludin, VE-cadherin, PECAM-1/CD31) from E65 onwards until P90. The expression profile was similar in visual and somatosensory cortex. Together, we report a rapid maturation of the vascular system in pig cortex. Regarding activity-dependent aspects, the data suggest that angiogenesis might be influenced by spontaneous rather than sensory-evoked activity.

**Author contribution:** Conceptualization and methodology: ES and PW; investigation and analysis: ES, KC; provided resources: CB, CB, MM, DU; visualization: ES, MM, DU; writing—original draft: ES, PW; writing—review and editing: ES, GM, PW; supervision and project administration: PW. All authors approved the manuscript.

**Summary statement:** Vascular impairment is associated with neurodegenerative disorders. As a groundwork for future studies, we quantitatively analyzed development of cortical vasculature in the emerging translational model, the pig.

**ETHICAL APPROVAL:** All applicable international, national, and/or institutional guidelines for the care and use of animals were followed. All procedures were in accordance with the ethical standards or practice of the institution at which the studies were conducted.

## INTRODUCTION

Most of the research on cortical development to date is still done with small rodents besides non-human primates and human. The pig as an alternative experimental and translational model is a precocial nest fledgling with almost fully formed sensory and motor systems at birth. The pig brain resembles the human brain with respect to its high cortical gyration index and neuron count (Zilles et al., 2013). A recent genome exploration of the pig (Teng et al. (2024) also demonstrates the high similarity of gene expression in various body organs between pig and human.

The vascular system delivers oxygen as well as hormones, glucose and metabolites, serves the tissue homeostasis and as waste disposal (Andreone et al., 2015). The vascular network is adapted to the local metabolic demand (Ji et al., 2021). Delivery of immune cells seeding the diploë of the skull and the meninges installs a line of defense which can rapidly contribute to protection against pathogens and infection. Interactions between endothelial cells and the neuronal environment are necessary for the formation of the blood-brain barrier (BBB) (Stewart and Wiley, 1981) which separates vasculature and CNS. The cellular elements contributing to the BBB are summarized under the term neurovascular unit. Well characterized are the roles of astrocytes and microglia (Thurgur and Pinteaux, 2019). Furthermore, pericytes participate in the control of blood flow (Armulik et al., 2010; Attwell et al., 2016; Grubb et al., 2020; Hall et al., 2014). Both, astrocytes and capillary pericytes were observed to be innervated by local neurons for neurovascular coupling (Andreone et al., 2015; Kisler et al., 2017b). A permanent crosstalk between the contributing cell types ensures the functional integrity of the BBB in the healthy brain, or the recovery after a pathological insult (Kozma et al., 2021; Liu et al., 2020).

Recently, neurovascular unit disruption has been studied in newborn pig with fetal growth restriction. Treatment with the anti-inflammatory drug ibuprofen attenuates BBB disruption, reduces overall pro-inflammatory responses, and promotes tighter vasculature-astrocytic end feet contacts which restores neurovascular unit integrity (Chand et al., 2022). In the case of neurological diseases such as Alzheimer’s disease (Clark et al., 2022; Kisler et al., 2017a), multiple sclerosis (Kaushik et al., 2021), or schizophrenia (Greene et al., 2017) the crosstalk can be impaired. This can lead to an altered BBB permeability enabling the infiltration of toxins, pro-inflammatory molecules, and cells into the brain (Cayrol et al., 2008; Haruwaka et al., 2019). Developmentally, the absence of pericytes during embryogenesis leads to damaged integrity and leaky BBB (Daneman et al., 2010; Mäe et al., 2021). A number of recent discoveries have extended our knowledge. A fourth meningeal layer, the subarachnoid lymphatic-like membrane has been characterized (Møllgård et al., 2023) with the hypothetical function of a filter membrane. Additionally, evidence emerged that the dura mater is directly connected to the brain via bridging veins that form structures termed arachnoid cuffs (Smyth et al., 2024). Moreover, vascular endothelial cells collected from different brain regions display a surprising variability in gene expression, which might even influence progression rates of Alzheimer’s disease (Bryant et al., 2023).

The vascular system develops in synchrony with the brain (Segarra et al., 2019). In the chicken embryo, the neural tube is already encircled by a vascular plexus which subsequently extends endothelial sprouts penetrating into the neural tissue (Feeney and Watterson, 1946). Cajal-Retzius neurons (Meyer, 2010) promote endothelial cell proliferation and angiogenesis (the creation of new vessels from preexisting ones), which stimulates radial glia cell attachment to the gliovascular basal lamina to ensure proper cortical neuron migration (Biswas et al., 2020). Radial glia cells also anchor to local blood vessels and disruption of these interactions impairs the immigration of interneurons (Tan et al., 2016). Depleting vascular reelin signaling alters neocortical vascularization and disrupts the assembly of radial glial cell contacts at the pial surface. Thus, abrogation of vascular reelin signaling partially recapitulates the neural defects observed in the absence of reelin protein (Thomas, 2018). Later on, neuronal activity influences vascular refinement as has been shown for the laminar-specific angiogenesis in visual cortex of marmoset (Fonta and Imbert, 2002).

In previous studies of fetal pig cortex, we describe the rapid development of the Neuropeptide Y neuron system (Ernst et al., 2018; Sobierajski et al., 2022)(Ernst et al., 2018) and of microglia in the cortical laminar compartments (Sobierajski et al., 2022). Also, myelination begins early with the first oligodendrocyte progenitors found at E45 (gestation in pig is 114 days) and large amounts of myelinated axons being present already at E100. Expression of myelin-related proteins commences 1-2 weeks earlier in the somatosensory compared to the visual cortex which suggests a driving role of specific somatosensory activity (Sobierajski et al., 2023). The present study analyzes vascular development in visual cortex, and the expression of selected proteins in visual and somatosensory cortex.

## 2. MATERIAL AND METHODS

### 2.1 Animal material

The material for the immunohistochemical staining is from our fetal pig brain collection (Ernst et al., 2018; Sobierajski et al., 2023). Our model of choice is a non-domesticated form, the European wild boar, *Sus scrofa* (Linné, 1758; Ritter et al., 2023). One argument for the feral representative comes from the fact that domestication of pig results in a substantial reduction of brain weight by about 41% in comparison to wild boar (Böndel, 2017). The second reason is that the material was obtained at no extra costs from the Üfter Mark area managed by the Regionalverband Ruhr Grün. Fetuses derived from mostly young (first pregnancy) sows and piglets individually hunted for population control in accordance with the German Game Law (Jagdrecht) or killed in road accidents. Law requests disposal of viscera including sexual organs. During evisceration at the Forsthof Üfter Mark, the uteri were examined for pregnancies. Embryonic membranes were removed, and fetuses immersed in cold 4% paraformaldehyde (PFA) in 0.1M phosphate buffer pH 7.4. The P5 domestic “German Landrace” piglet was donated by the Institutes of Physiology and Anatomy, Medical Faculty, University Mannheim (donated by Prof. Martin Schmelz and Prof. Dr. Maren Engelhardt). Staging has been done using the crown-rump-length formula (Henry, 1968) for European wild boar and by external features. “Gestation” is the proportional age with E114 (birth) set to 1; it has been calculated to compare e.g. to data in fetal sheep. Fine dissection of the brain and the body organs was done after transport to Ruhr University. The meninges were largely removed to minimize cryostat cuttting artifacts.

### 2.2 Tissue processing and immunostaining procedures

All litters prepared for immunohistochemistry and for protein blots, respectively, have been listed in detail earlier (Sobierajski et al., 2023; see Table 1). Dissected brains were immersion-fixed in 4% paraformaldehyde in 0.1 M phosphate buffer pH 7.4 with 5% (vol/vol) water-saturated picric acid for about 2 weeks at 8°C (refreshed once). After cryoprotection, tissue slabs were stored frozen in TissueTek (Sakura Finetek, Alphen aan den Rijn, Netherlands) at −80°C until cutting. Coronal 25 µm cryostat sections of occipital cortex mounted on silanized slides were submitted to antigen retrieval (40 min citrate buffer pH 5-6 at 80°C and cooling down slowly for about 1 h) followed by immunohistochemistry as previously described (Sobierajski et al., 2022). Blood vessels have been stained with tomato lectin, isolectin B4 (IB4), or PECAM1. Tomato lectin is a superior marker for blood vessels in normal tissue (Battistella et al., 2021). IB4 marks vessels (Battistella et al., 2021; Gama Sosa et al., 2021). It also marks macrophages and activated microglia by binding to the Ret receptor (Boscia et al., 2013). IB4 labeling intensity decreases from fetal to postnatal in rodent (Wu et al., 1994). Antibodies and lectins are listed in Table 1. Immunofluorescent labelling was combined with a quenching of autofluorescence by incubation in Sudan Black B (0.2% w/v dissolved in 70% isopropyl alcohol) for about 60 min before incubation of the secondary antibody. A brief incubation with DAPI was done before coverslipping to delineate cortical layers and mark cell nuclei. Alternatively, we used biotinylated secondaries followed by avidin-biotin-horseradish peroxidase complex and 3,3-diaminobenzidine. The DAB reaction product was intensified with 1% OsO_4_ in phosphate buffer for 1-2 minutes. Finally, the sections were dehydrated and coverslipped with DPX (Sigma Aldrich, Steinheim, Germany). In some cases, a counterstain with thionin was done before dehydration. No specific staining remained after omitting the primary and/or secondary antibodies except for autofluorescence of blood cells.

**Table 1.** Antibodies and reagents for immunohistochemical staining and protein blotting. IHC, immunohistochemistry with DAB staining; IF, immunofluorescence; WB, Western blot.

| Primary antibodies | Species, label, method | Source, order number, RRID | Dilution |
| --- | --- | --- | --- |
| Alpha-smooth muscle actin | Rabbit, IF, WB | Abcam, Cambridge, UK, Cat# ab5694, RRID: AB_2223021 | 1:200 (IF)<br>1:1000 (WB) |
| Aquaporin 4 | Guinea pig, IF, WB | Synaptic Systems, Göttingen, DE, Cat# 429004, RRID: AB_2802156 | 1:800 (IF)<br>1:1000 (WB) |
| Claudin-5 | Mouse, IF, WB | ThermoFisher Scientific, Darmstadt, DE, Cat# 35-2500, RRID: AB_2533200 | 1:200 (IF) |
| Glial fibrillary acidic protein | Rabbit, IF | Dako A/S, Glostrup, DEN, Cat# Z0334, RRID: AB_10013382 | 1:400 (IF) |
| Isolectin B4 (*Bandeiraea simplicifolia*) | FITC, IF | Enzo Life Sciences GmbH, Lörrach, DE, Cat# ALX-650-001F | 1:400 (IF) |
| Occludin | Guinea pig, WB | Synaptic Systems, Göttingen, DE, Cat# 447005, RRID: AB_2884909 | 1:1000 (WB) |
| Platelet-derived growth factor receptor beta | Goat, IF, WB | R&D Systems, Minneapolis, MN, USA, Cat# AF385, RRID: AB_355339 | 1:500 (IF)<br>1:1000 (WB) |
| Platelet endothelial cell adhesion molecule 1 | Guinea pig, IF, WB | Synaptic Systems, Göttingen, DE Cat# HS351004, RRID: AB_2620105 | 1:300 (IF) |
| Platelet endothelial cell adhesion molecule 1 | Mouse, IF, WB | Santa Cruz Biotechnology, Dallas, Texas, USA, Cat# sc-376764, RRID: AB_2801330 | 1:1000 (WB) |
| Vascular endothelial cadherin | Goat, WB | Santa Cruz Biotechnology, Dallas, Texas, USA, Cat# sc-6458, RRID: AB_2077955 | 1:1000 (WB) |
| β-actin (housekeeping protein) | Mouse, WB | Sigma-Aldrich, St. Louis, MO, USA, Cat# A1978, RRID: AB_476692 | 1:6000 (WB) |
| β-Tubulin (housekeeping protein) | Mouse, WB | Sigma-Aldrich, St. Louis, MO, USA, Cat# T8660, RRID: AB_477590 | 1:3000 (WB) |
| Tomato lectin (*Lycopersicon esculentum*) | biotin, IF, IHC | Sigma-Aldrich, St. Louis, MO, USA, Cat# L0651 | 1:500 (IF, IHC) |
| **Secondary antibodies** |  |  |  |
| Streptavidin | DyLight™ 594, IF | ThermoFisher Scientific, Darmstadt, DE, Cat# 21842 RRID: AB_2619631 | 1:1000 (IF) |
| Anti-rabbit | Donkey, Alexa-488, IF | Thermo Scientific, Waltham MA, USA, RRID: AB_2687506 | 1:1000 (IF) |
| Anti-mouse | Goat, Alexa-568, IF | Invitrogen, Carlsbad, CA, USA, RRID: AB_2534013 | 1:1000 (IF) |
| Anti-guinea pig | Goat, Alexa-488, IF | Thermofisher Scientific, Darmstadt, DE, Cat# A11073, RRID: AB_2534117 | 1:1000 (IF) |
| Anti-guinea pig | Donkey, Alexa-594, IF | Dianova, Cat# 706-757-148 | 1:1000 (IF) |
| Anti-guinea pig | Goat, Star Green, IF | Abberior dyes, Göttingen, DE, Cat# STGREEN-1006-500UG | 1:200 (IF) |
| Anti-rabbit or anti-mouse | Goat, Star Red, IF | Abberior dyes, Göttingen, DE, Cat# STRED-1002-500UG | 1:200 (IF) |
| Anti-rabbit or anti-mouse | Goat, Star Orange, IF | Abberior dyes, Göttingen, DE, Cat# STORANGE-1001-500UG | 1:200 (IF) |
| Anti-mouse | Rabbit, alkaline phosphatase, WB | Dako, Hamburg, DE, Cat# D0314 | 1:5000 (WB) |
| Anti-goat | Donkey, alkaline phosphatase, WB | Life Technologies, Carlsbad, CA, USA, Cat# A16002, RRID: AB_2534676 | 1:2000 (WB) |
| Anti-rabbit | Goat, alkaline phosphatase, WB | Dako, Hamburg, DE, Cat# D0487, RRID: AB_2617144 | 1:3000 (WB) |
| Anti-guinea pig | Goat, alkaline phosphatase, WB | ThermoFisher Scientific, Darmstadt, DE, Cat# A18772 | 1:5000 (WB) |
| **Reagents** |  |  |  |
| Tetrahydrofuran | Lipid removal | Sigma-Aldrich St. Louis, MO, USA, Cat# 186562 | 50% (v/v) |
| Nycodenz | Transparent rendering | Serumwerk Bernburg AG, Bernburg, DE, Cat#18003 | 80% (w/v) |
| DAPI | Fluorescent counterstain | Sigma-Aldrich, St.Louis, MO, USA, Cat# D9542 | 1:2500 |
| Sudan Black | To quench autofluorescence | Merck, Darmstadt, DE, Cat# 1387 | 0.2% (w/v) |
| ABC reagent | Horseradish peroxidase; | Vector Laboratories, Inc., Burlingame, CA, USA, Cat# PK-6100, RRID: AB_2336827 | 6 µl vector A/B per mL TBS |
| Diaminobenzidine DAB | Chromogen | Sigma Aldrich, Steinheim, DE, Cat# D12384 | 0.02% (v/v) |
| Osmium tetroxide | Intensification | Sigma Aldrich, Steinheim, DE, Cat# O5500 | 1% |

### 2.3 Tissue clearing

Vibratome sections with a thickness of 300 µm were treated with the “EZ Clear” method (Hsu et al., 2022). In brief, the slices were washed in tetrahydrofuran for 16 h in light-protected glass scintillation vials, then washed 4 times for 1 h each in distilled water. After washing, immunofluorescent labeling was performed with the blocking step increased to 1 day and the incubation times of the primary and secondary antibody each increased to 2 days at room temperature. Thereafter, sections were incubated for 1 day in EZ clear solution (7 M urea, 0.05% N-azide, 80% Nycodenz in 0.02 M phosphate buffer pH 7.4) until they rendered transparent. For confocal images, the slices were embedded with EZ view solution in µ-dishes (ibidi GmbH, Gräfeling, Germany) and flattened with a piece of foil to obtain a planar focus level.

### 2.4 Western Blots

As described (Sobierajski et al., 2023), tissue blocks of visual and somatosensory cortex were taken during preparation of the fetuses (ages E65, E80, E95, E100 and P90) and frozen on dry ice for storage. Small blocks were first pulverized on dry ice. Microspoon-sized samples of the powder were homogenized in standard RIPA buffer. Protein amount was determined with the Markwell protein assay. For SDS-PAGE, proteins were separated on 10% or 14% polyacrylamide gels (30 mg per lane) with a current voltage of 7 mA per gel overnight. Proteins were transferred to nitrocellulose membranes (BA 85, Schleicher and Schuell, Germany) with a tank blotter (Hoefer, San Francisco, CA, USA). Smaller proteins were transferred with buffer containing 20% MeOH, proteins above 100 kDa were transferred with buffer containing 15% MeOH and 0.05% SDS. Membranes were cut into horizontal stripes (Engelhardt et al., 2018) containing the desired protein bands such that up to 8 proteins of different molecular weights could be concurrently detected in each lysate. Membranes were blocked with TBS (50 mM Tris-HCl, pH 7.4, 150 mM NaCl) containing 5% bovine serum albumin for 2 h. Primary antibodies (Table 1) were incubated overnight at 4°C, followed by washes for 10 min in TBST (50 mM Tris, pH 7.4, 150 mM NaCl, 0.1% Tween-20) and 10 min in TBS. Appropriate alkaline phosphatase-conjugated secondaries were incubated for 90 minutes. Blots were stained with 0.18 mg/mL BCIP and 0.35 mg/mL NBT in an alkaline buffer (100 mM Tris-HCl, pH 9.5, 50 mM MgCl_2_), rinsed with water and dried.

### 2.5 Analysis

DAB-stained material was analyzed with light microscopy. The E45 plot was done from thionin-stained sections with the Neurolucida (MicroBrightField, Inc., Williston, Vermont, USA). Bright field photomicrographs were taken with a Zeiss Axiophot equipped with a CCD camera (PCO, Kelheim, Germany). Images and plots were arranged with Adobe Photoshop® (CS6 Extended, Version 13.0 x64) and Inkscape (open-source software). Regions of interest were selected with the aim to document the distribution of pericytes or vasculature in the laminar compartments at selected fetal and postnatal stages. Fluorescent images and tile scans were done with a Leica TSC SP5 confocal microscope (10x and 40× objective with 1.1 NA, 1024 × 1024 px). Global whole-picture contrast, brightness, color intensity and saturation settings were adjusted with Adobe Photoshop®. Scale bars were generated with ImageJ (MacBiophotonics) and inserted with Adobe Photoshop®.

STED imaging was performed on a commercial STED microscope (Facility Line, Abberior-Instruments, Germany) working at a repetition rate of 40 Mhz. Samples were immunostained using a FITC-coupled lectin or antibodies with appropriate secondaries conjugated to the dyes STAR RED, STAR ORANGE, and STAR GREEN as well as Alexa 568. STAR RED was imaged with excitation at a wavelength of 640 nm and time-gated fluorescence detection between 650-760 nm. STAR ORANGE and Alexa 568 were excited at 561 nm with time-gated detection between 571-630 nm for STAR ORANGE and 573-683 nm for Alexa 568. STAR GREEN and FITC were excited at 488 nm with time-gated detection between 498-551 nm. The Abberior MATRIX array detector was used to reduce unspecific background signal. The STED lasers had a wavelength of 775 nm and a pulse width of roughly 500 ps. An oil-immersion objective was used (UPLXAPO 60XO, NA: 1.42, Olympus, Japan).

For assessment of PDFGRβ+ cells to total cells in confocal images, regions of interest were arbitrarily placed over the selected cortical compartments. Labeled cells with a DAPI-positive nucleus in the optical plane were individually marked in the “3D-environment” function of Neurolucida 360 software and calculated per mm^2^ and per mm length of blood vessels, respectively. Data management was done in Microsoft Excel 365. All graphs were made with SigmaPlot 12.3 (Systat Software GmbH).

The reconstruction and evaluation of blood vessel data was done with the Imaris 10.1. software (Oxford Instruments). Regions of interest were arbitrarily placed over the selected cortical compartments. Z-stacks were mostly in the range of 100-200 µm, limited either by microscopic range, computing power, and, in few cases, depth-depending fading of antibody staining. First, a semi-automatic layout of the 3D vessel pattern was generated with the “filament” algorithm. Next, manual corrections were done such as deleting redundant or adding missing parts and branch points, and correcting vessel thickness. The two litters aged E68 and E72 were combined under E70, E100 data were pooled with data from E110 because the two were not different. Statistical measures were taken from Imaris, data management was done as described above. A major technical consideration concerns the automated data acquisition from digital images promized by the software tools. In our case the tools deliver a coarse layout of the vessel pattern but that requires a serious workover by trained examiners because e. g. thin vessel connections would have been missed. A similar note towards involving the best hightech available, the human retina and brain, has been given for working with automated serial reconstructions of EM images (Shapson-Coe et al., 2024; Wu et al., 2024).

Protein blots were photographed, and relative band intensities were determined with ImageJ. All protein bands were normalized to the 42 kDa β-actin or the 55 kDa β-tubulin band, which served as control for loading equivalency. Values were normalized to the P90, which was considered adult. For every protein, at least three independent lysates from cortex of two animals (mostly from the same litter) were tested for every age.

## RESULTS

### The early phase of angiogenesis

Stage E45 is shortly after the transition from embryonic to fetal in pig, the hippocampal *anlage* is recognizable, and the stage is comparable to 15^th^/16^th^ gestational week in human (Meyer, 2010). Examination of occipital cortex and midbrain (Fig. 1 A) revealed a dense and very uniform network of vasculature in the midbrain *anlage* (Fig. 1 B) whereas only very few vessels can be seen in the cortical *anlage* (Fig. 1 C-D). Large and small diameter vessels were present in the meninges (Fig. 1 C, inset C1), and vessels are innervated by Neuropeptide Y-positive fibers (Ernst et al., 2018). At E45, gyrification had not yet begun and the transient cortical layers such as cortical plate (CP) and subplate (SP) were narrow in depth. Intermediate zone (IZ) and ventricular zone (VZ) occupied about two thirds of the cortical depth (Fig. 1 C, inset of a thionin-stained panel) and most of the vessel elements were within the proliferative VZ/SVZ presumably to supply the stem cell niche. Vessels within the outer layers were mostly unbranched, while those in VZ/SVZ had already developed some branches and anastomoses (Fig. 1 D, inset D1). No preferred orientation of the vessels could be recognized. In the IZ, as well as the SP, CP, and marginal zone (MZ), the vessel density was low. This qualitative observation suggests a high angiogenic activity at the begin of fetal development. The substantially advanced microvessel pattern in the midbrain matched well with the finding of numerous microglia cells and oligodendrocyte progenitors reported earlier for the midbrain *anlage* (Sobierajski et al., 2022; Sobierajski et al., 2023).

**Figure 1.**
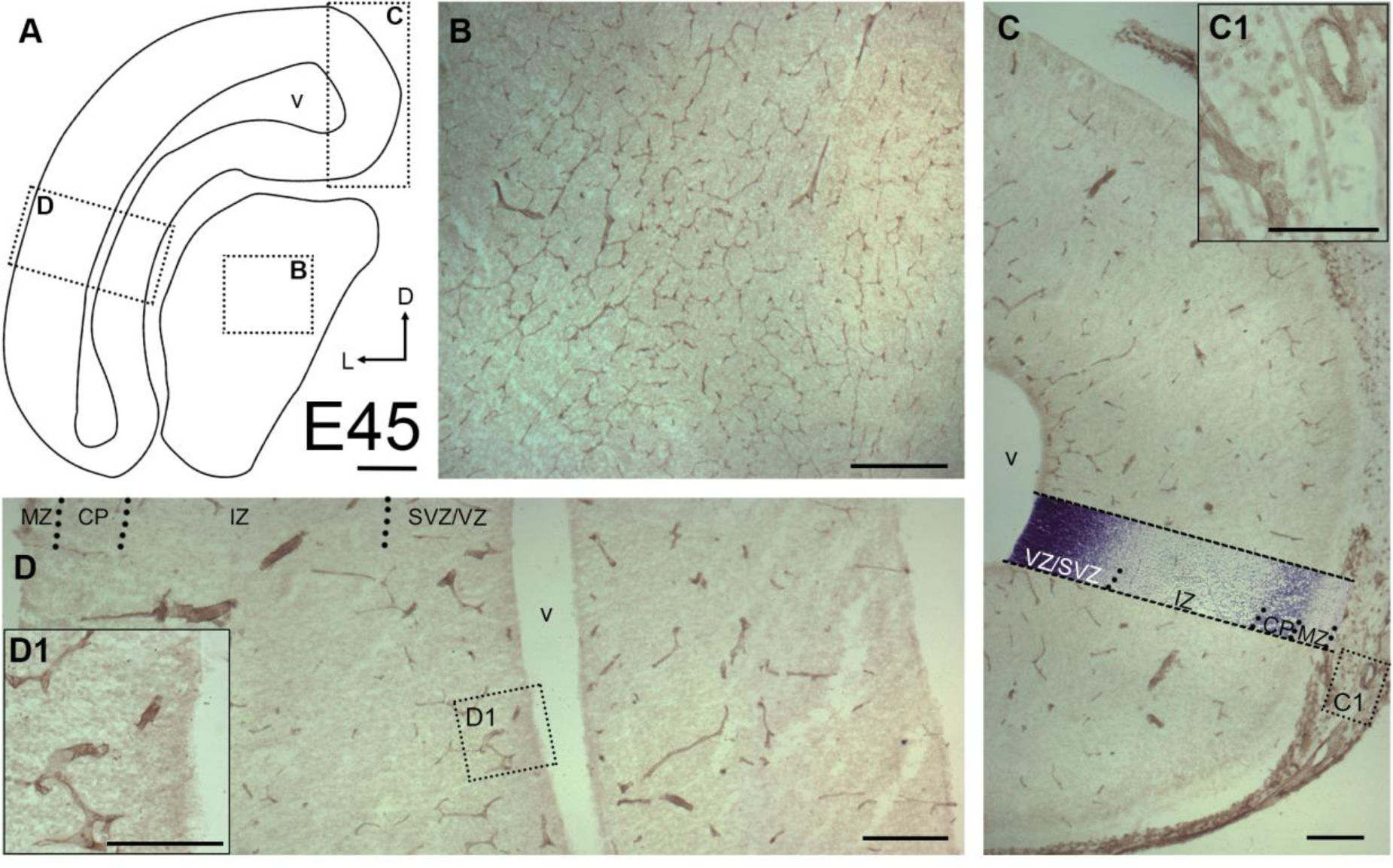
Qualitative assessment of blood vessels at E45. A. Outline of a section through occipital cortex and midbrain *anlage*. Arrows: D, dorsal; L, lateral. Boxed regions are shown at higher magnification in the insets. B. The midbrain a*nlage* already displayed a dense vascular network. C. Cortex, medial aspect. Meninges have been partially left attached to avoid damaging of the cortex. C1. The meninges displayed a dense network of thick and thin vessels. A thionin-staining panel has been overlayed to demonstrate the laminar compartments. D. Cortex, lateral aspect. The developing upper layers display hardly any vessels while VZ/SVZ display more vessels. Abbreviations apply to all following figures: MZ, marginal zone; CP, cortical plate; IZ, intermediate zone; SVZ, subventricular zone; VZ, ventricular zone; v, ventricle. Scale bars: 200 µm in A-D; 100 µm in D1.

### Midfetal to postnatal stages

Next, high-volume images of selected gyri of the visual cortex at E70 to P90 were imaged (Fig. 2). A massive increase in vascular density has occurred compared to E45. Given the enormous expansion of the pig brain and cortex (Ernst et al., 2018), we selected a major gyrus at E70 and successively smaller gyri at E85 and E100 in order to show the patterns roughly at the same magnification. As an example of the laminar expansion, the thickness of MZ/layer 1 (L1) increased dramatically from E70 to E100 (from 123 µm to 385 µm; measured midlevel at gyral flanks, see Fig. 2 A1 and C1). The gray matter (GM) thickness (border of MZ/L1 to border of white matter (WM)) measured at the same position increased from ∼540 µm at E70 to 890 µm at E100. At E70, long, unbranched vessels extended from deeper WM towards the apex of the gyri (Fig. 2 A). Thinner branches bended off and extended through the GM layers, and connected to vessels which penetrate the MZ/L1 from the pial surface. All vessels formed thinner branches on their way through the cortical layers, which are commonly referred to as penetrating arterioles. GM layers, in particular the cell-dense CP, were only sparsely covered by vessels.

**Figure 2.**
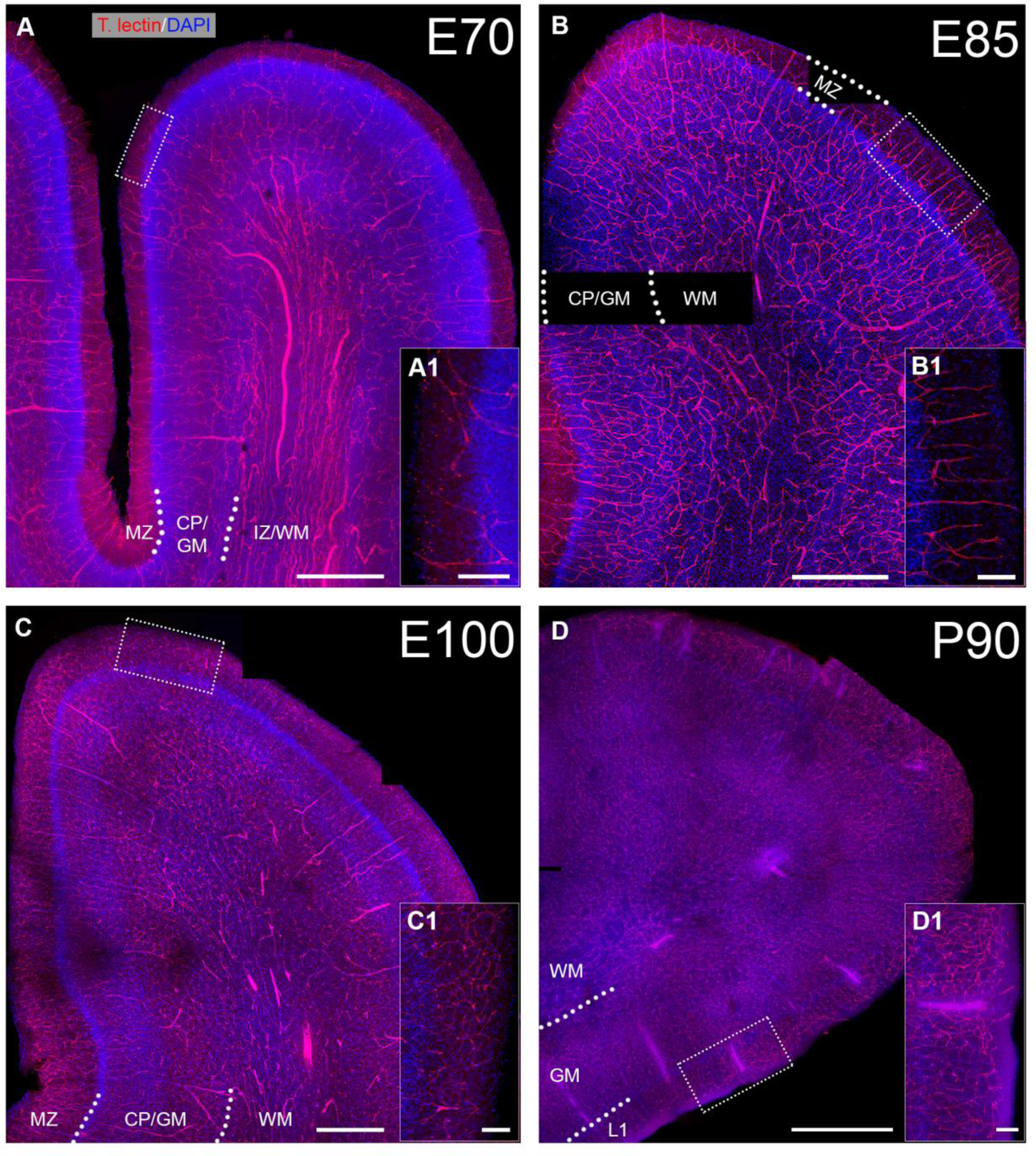
Blood vessel pattern from E70 to P90. Cleared brain sections of ∼300 µm thickness were stained with Tomato lectin. Selected gyri of visual cortex were imaged with tile scans. A. E70, note the long unbranched vessels with large diameter in the IZ/WM. B. E85, larger numbers of perpendicular vessels can be seen that branch in the CP/GM. C. E100. D. P90. A1, B1, C1, D1. For every age, a magnification of apical MZ/L1 is shown to compare the thickness of L1 as well as vascular density. Scale bars: 500 µm in A-C, and 1000 µm in D; 100 µm in A1, B1, C1, D1.

At E85, vessel density had increased forming a seemingly homogeneous network in all layers of the GM (Fig. 2 B) with higher numbers of penetrating arterioles. The pattern at E100 differed in that the density of finer capillaries had increased in GM and L1 accompanied by an increased density of penetrating arterioles (Fig. 2 C). The vascular density of deeper layers and WM had changed less compared to E85. At P90 (Fig. 2 D), the vessel density had again increased visibly as compared to E100 suggesting that the angiogenic activity remained high after birth.

### Development of the vascular network in the cortical compartments

During embryonic development, the vascularization of the telencephalon is driven by two plexi, the periventricular vascular plexus and the perineural vascular plexus which is wrapped around the developing CNS (Kaushik et al., 2020; Kurz et al., 1996; Penna et al., 2021). The perineural vascular plexus runs along the pia mater and grows specialized “tip” cells which enter the MZ/L1 perpendicularly. This way regularly spaced vertical vessel trunks emerge. Tip cells direct the sprouting vessel very similar to how growth cones steer elongating neurites. Proliferation of endothelial cells delivers the material for the vessels and the penetrating arterioles, and finally capillaries.

For a quantitative assessment, a linear measure of the surface has been taken at arbitrarily selected positions. The number of vertical vessel trunks was determined (Fig. 3 A-C). Furthermore, number of vessels crossing the border of MZ/L1 to CP/GM was counted as a readout for the branching within the MZ (Fig. 3 D). On average, at E70, 15 penetrating vessels per 1 mm surface were counted at the pial surface, and 24 vessels/mm were counted at the CP/GM border. At E85, 19 penetrating vessels/mm at the pial surface, and 21 vessels/mm were counted at the CP/GM border. At E100, 27 penetrating vessels/mm were counted at the pial surface, and 39 vessels/mm were counted at the CP/GM border. Together, within a period of 30 days the number of penetrating vessels doubled and many appear to branch within the MZ/L1 concurrent with the substantial increase in brain volume (Ernst et al., 2018).

**Figure 3.**
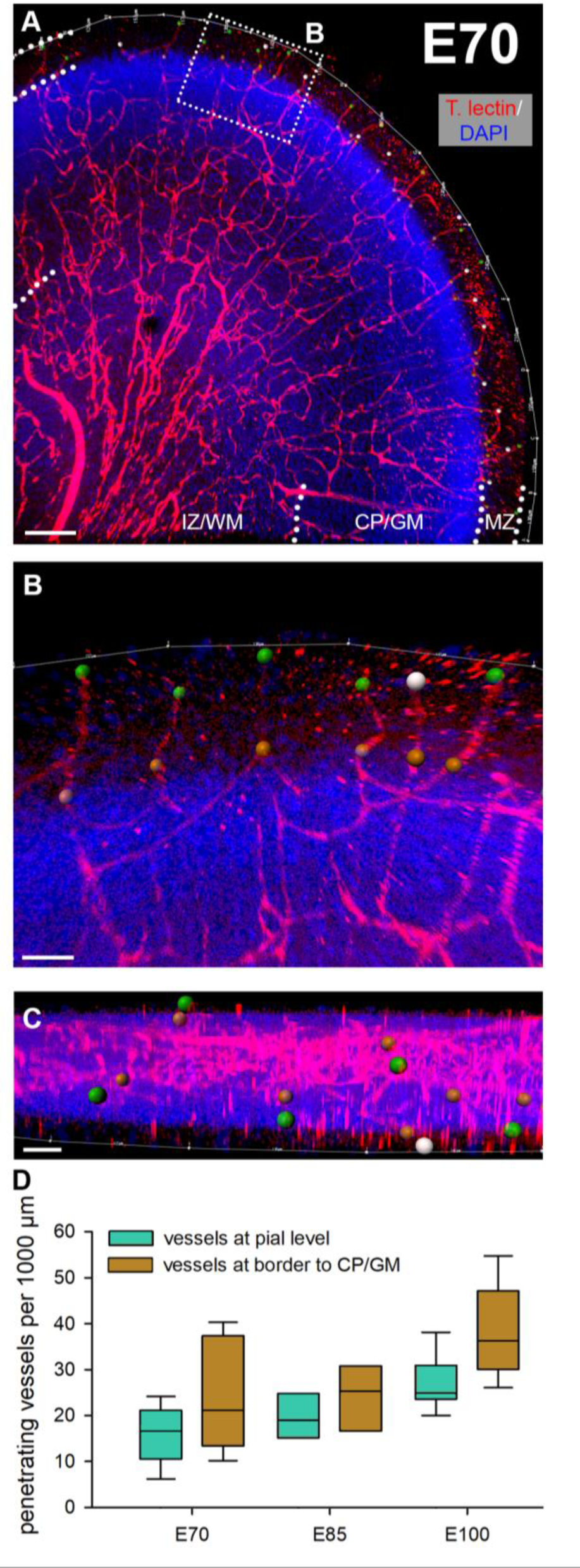
Penetrating blood vessels in visual cortex. From E70 to E100, the number of penetrating vessels that enter from the pial surface were plotted. A. Tomato lectin-stained gyrus at E70. The dotted rectangle is magnified in B. B. The pial surface length was outlined (thin white line) and measured. Green dots represent vessels at their entrance positions into the cortex. Gray dots represent vessels for which the entrance point could not exactly be determined due to leaving the z-axis of the confocal stack (at least 200 µm depth were imaged). Yet, it was reasonable to assume they might have also entered from the pial surface. Brown dots mark vessels that cross the MZ/L1 to CP/GM boundary. C. Z-view of the region of interest shown in B. D. Quantitative assessment of penetrating vessels at pial level and at level of MZ/L1 to CP/GM border. E. Measurement of the nearest neighbor distances of vessels that entered the cortex from the pial surface. Scale bars: 150 µm in A; 40 µm in B; 80 µm in C.

Next, a quantitative assessment of vessel densities in cortical layers was done to assess the progression of angiogenesis. For this purpose, cleared slices of approximately 300 µm thickness were stained with immunofluorescence, imaged, and analyzed with the Imaris software (workflow is shown in Fig. 4 A). The ventricular zone is a transient cell layer. It decreased substantially between E100 and P30 in pig and was therefore excluded at P30. Source data of all assessed parameters are summarized in the Excel Table S1.

**Figure 4.**
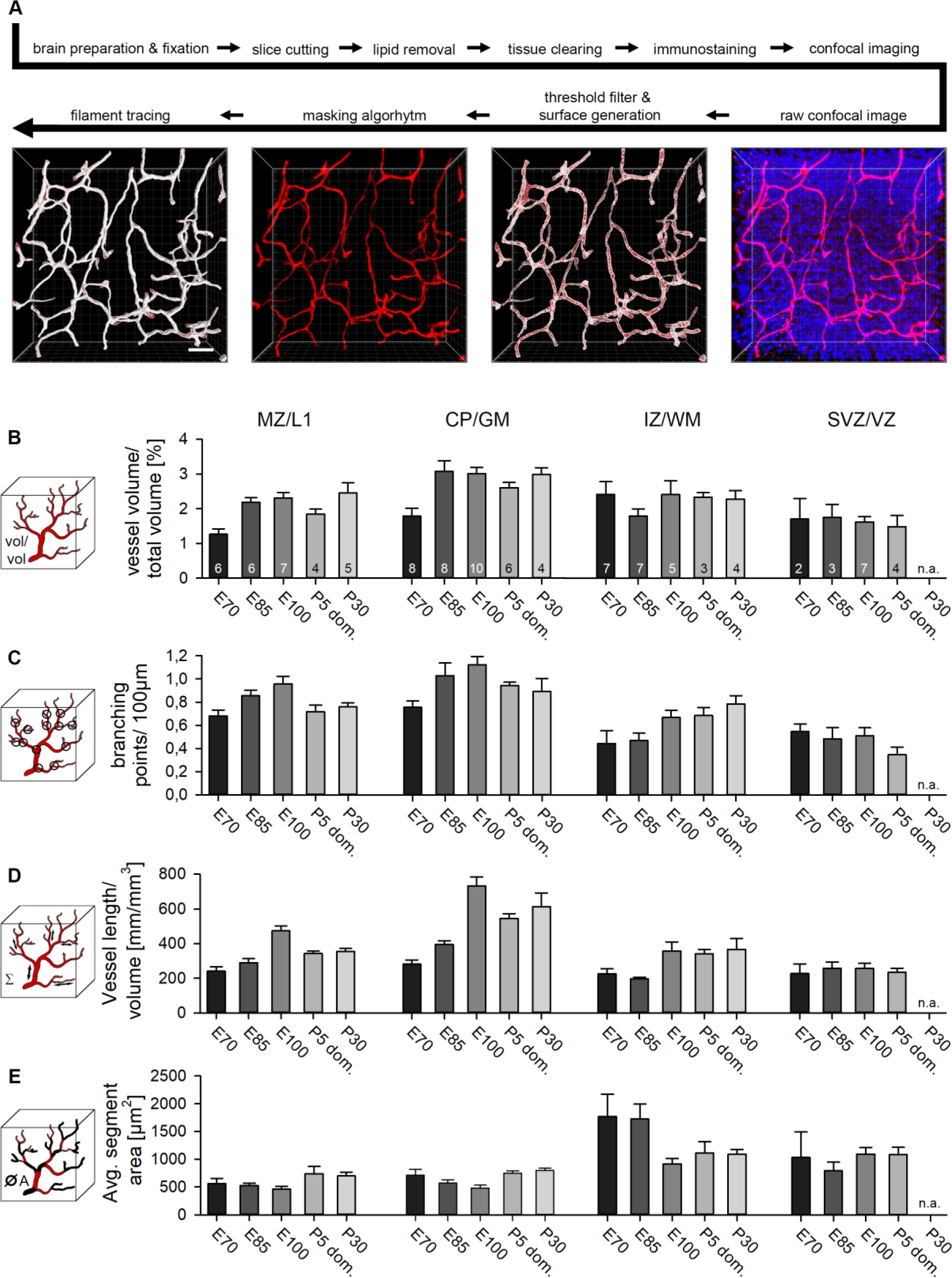
Quantitative assessment of vasculature. A. Experimental workflow from dissection to filament tracing (Imaris). The raw confocal image was processed with a threshold filter and a masking algorithm. Next, the “surface” tool was used to create a surface for volume measurement and masking algorithm. The “filament” tool yielded a semiautomatic layout of the vessel pattern which has in all cases been manually refined. Vascular parameters were adressed in four laminar compartments. If present, the SP was included in the IZ/WM compartment. B. Volume of all vessels compared to the total sample volume. C. Vessel branching points were calculated per 100 µm. D. The length of vessels was set in relation to the total sample volume. E. The average vessel segment area. Vessel sections between two branching points were defined as segments. The numbers in the bars represent the number of non-adjacent regions of interest that were assessed for this compartment. At P30, no sufficient SVZ/VZ area could be found suitable for reliable reconstruction of that compartment; n. a., not applicable. Error bars represent the mean ± S.E.M.. Scale bars: 40 µm in A.

First, we measured the volume occupied by blood vessels in relation to the total volume of the sample (Fig. 4 B). Vessels gradually obtained more total space in GM layers while vessel volumes in SVZ/VZ remained comparatively low throughout development. A maximum of ∼3.0% vessel volume of total sample volume was reached in the CP/GM around E100 and remained until P30. In the IZ/WM, vessel volume was at a stable level of ∼2.35% at E70 and stagnated until P30. Interestingly, at the border between VZ and ventricle, nearly no vessels could be found (Fig. S1 A). This zone has been described to harbor an avascular niche for the apical neural progenitor cells which reside in contact with tip cells of vessel sprouts (Komabayashi-Suzuki et al., 2019).

Interestingly, readouts for vessel volume, branching, and segment length of the domestic P5 were somewhat lower than the E100/110 in the MZ/L1 and mainly in CP/GM (Fig. 4 B, C, D). This might suggest that vascular development in domestic pig tends to proceed slower or to remain below the measures reached by wild boar individuals.

In addition, the average diameter of vessel segment was determined (Fig. S1 B). The largest vessel diameters were measured in the IZ/WM (4.88 - 6.71 µm) and SVZ/VZ (4.89 – 5.92 µm), followed by the CP/GM (3.58 - 5.11 µm) and MZ/L1 (3.03 – 4.87 µm) with smaller vessels on average. Vessel diameter did not change much in MZ and CP/GM from E70 to E100, and only subtly increased at postnatal age.

Second, the number of branching points per 100 µm vessel length was determined (Fig. 4 C). A maximum vessel branching seems to be reached in the gray matter shortly before birth with 1.18 branching points/100 µm at E100, then declining slightly and stabilizing postnatally. Overall lower branching was seen in IZ/WM and SVZ/VZ. Third, the total length of the reconstructed vessels per volume was measured in mm/mm^3^ (Fig. 4 D). The vessel length matches the results of the relative volume occupied by blood vessels and follows a similar development, as expected. Again, the CP/GM displayed the highest density with 733 mm/mm^3^ at E100. Likewise, the average vessel length in the SVZ/VZ was the lowest of all cortical layers.

Fourth, the average blood vessel segment area was determined (Fig. 4 E). A segment was defined as the part between two branching points. Both, in MZ/L1 and CP/GM, the segment area displayed only minor fluctuations. The largest change occurred in IZ/WM. Here, the average vessel area was initially 3-4-fold higher than in GM, only to decline substantially at E100 to a plateau persisting until P30. A similar picture can be observed in the data of the average segment volumes and segment lengths (Fig. S1 C, D). Compared to MZ/L1 and CP/GM, there were fewer vessels in the deeper cortical compartments WM and SVZ/VZ. This finding increased the relative influence of individual large vessels occasionally encountered in the arbitrarily selected ROI. It explains the overall higher S.E.M. of the statistical analysis. Also in human and rhesus monkey, the vessel density declines towards the WM (Lauwers et al., 2008; Weber et al., 2008).

### Establishment of blood-brain-barrier cellular components

The BBB becomes gradually assembled (Daneman et al., 2010). Here, we analyzed time of appearance of major structural and molecular components of the BBB (Fig. 5). Essential for barrier function are astrocytes whose endfeet are ensheathing capillary vessels contributing to the membrana limitans gliae perivascularis this way (Haug, 1971). At E45, the considerable differences described above (Fig. 1) between the midbrain *anlage* and the cortex can again be observed (Fig. 5 A-D). Astrocytic end feet immunopositive for the water transporter aquaporin 4 (AQP4) establishing contact with capillaries were present in the midbrain *anlage* (Fig. 5 A). This contrasted with the situation in the cortical VZ, where astrocytes cannot be found at E45 (Fig. 5 C), except for first signs of AQP4+ end feet detected close to the lateral ventricle (Fig. 5 D, small inset). Western blots confirmed the expression AQP4 protein at E65 (Fig. 7 A). As reported (Sobierajski et al., 2023), lysates have been made from the apex of gyri and thus are enriched for GM. Together, this suggested that vessel-contacting endfeet develop first in deep laminar compartments and later in GM. An intact BBB depends on functional tight junctions which act as physical seal between endothelial cells. Indeed, the tight-junction protein claudin-5 was detectable in capillaries at E45 in deeper cortical compartments suggestive of being the earliest evidence for BBB development.

**Figure 5.**
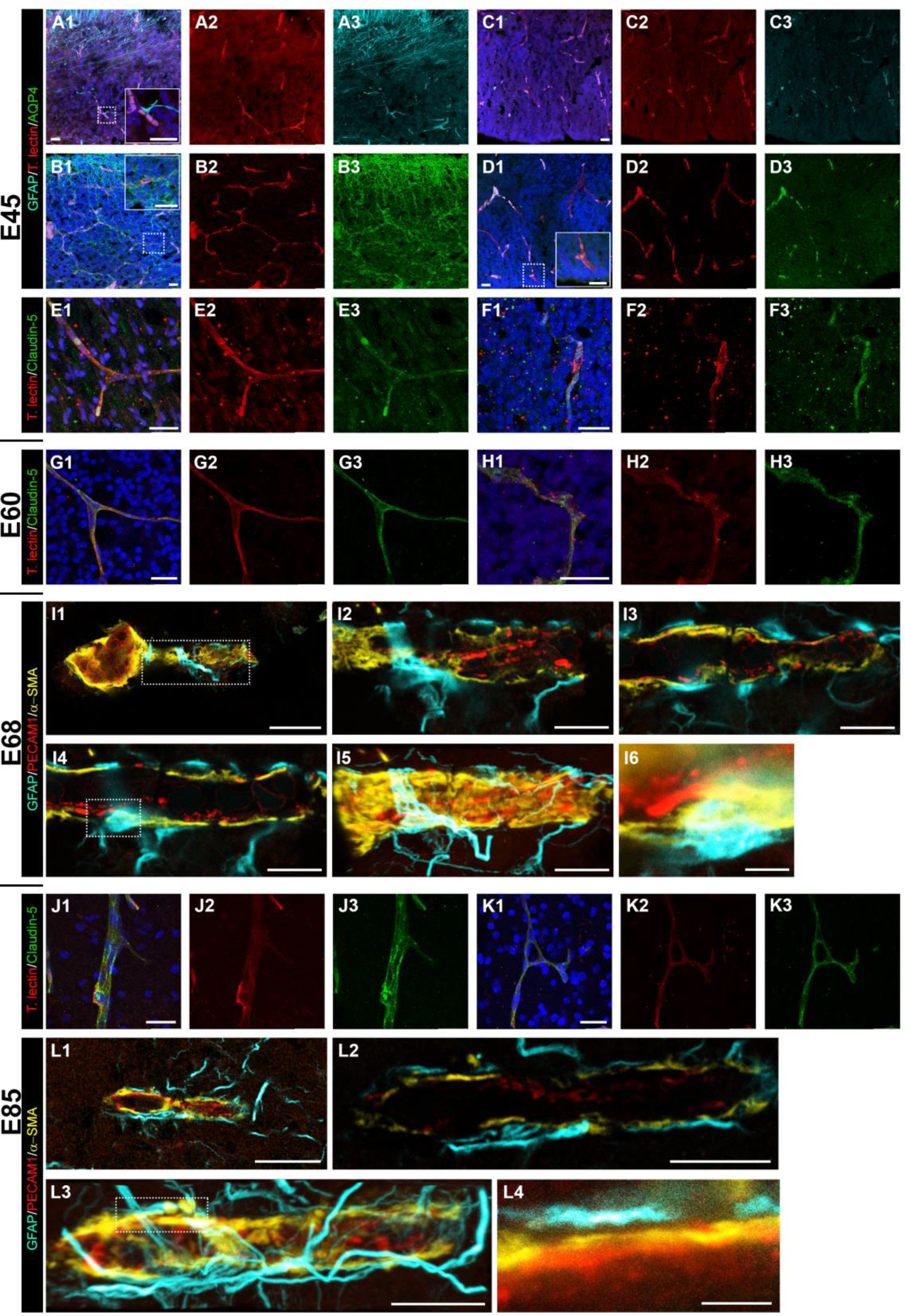
Establishment of the neurovascular unit. A, B. Midbrain *anlage* at E45. C-F. Cortex at E45. Shown are maximal projections, merged to the left, next to the single channels. A, C. GFAP staining. Note the presence of GFAP+ cell processes contacting blood vessels in midbrain (magnified in the inset). Cortical VZ is still void of GFAP+ cells or processes. B, D. AQP4+ staining. Note glial end feet building a barrier around capillaries in midbrain (magnified in the inset). The cortical VZ is nearly void of AQP4+ end feet. First indications are weakly stained puncta near capillaries close to the ventricle (magnified in the inset). E, F. Claudin-5 staining in cortex. E1-E3. Immunoreactive vessels in IZ/WM. F1-F3. Immunoreactive vessels in VZ. G, H. Cortex at E60; claudin-5 staining. G1-G3. Immunoreactive vessels in upper GM. H1-H3. Immunoreactive vessels in VZ. I. Cortex at E68; STED images. I1. An arteriole with branching capillary in the CP/GM. I2-I4. Single z-stack images at different levels of the vessel shown in boxed region of I1. I5. Maximum projection; note the layer of α-SMA and GFAP+ elements partially ensheathing the blood vessel. I6. Astrocytic cell body closely attached to the blood vessel; magnified from boxed region of I4. J, K. Cortex at E85. J1-J3. Claudin-5+ penetrating arteriole in CP/GM. K1-K3. Claudin-5+ vessel anastomosis in IZ/WM. L. Cortex at E85; STED images. L1. Thin capillary in the CP/GM. L2. Single z-stack image of vessel shown in L1. L3. Maximum projection. L4. Close-up of boxed area of L3. Note the gap between the astrocytic process and the surface of the blood vessel. Scale bars: 25 µm in A-H; 20 µm in I1 and L1; 10 µm in I2-I5 and L2-L3; 2 µm in I6 and L4.

At E60, claudin-5 immunoreactivity was present in all layers suggestive of an existence of stable tight junctions (Fig. 5 G1-G3, H1-H3). High-resolution STED images of vessels were taken at E68 (Fig. 5 I1). Here, we show an arteriole with a branching capillary in the cortical GM (Fig. 5 I2-6). We defined every blood vessel as a capillary that had a diameter below 8 µm, based on reports that first, the human capillary diameter underlies a Gaussian distribution with a mean of 6.23 ± 1.3 µm (Cassot et al., 2006) and second, around 80% of all vessels are capillaries (Bryant et al., 2023). Images of the individual z-layers show the assembly in detail. PECAM1 was mainly expressed on endothelial cells and particularly intensive at the contact points between neighboring cells. The actin isoform α-smooth muscle actin (α-SMA), is known to be expressed in arterioles and capillaries, but also in a significant number of mid-capillary pericytes mediating their contractility (Bandopadhyay et al., 2001). α-SMA forms a layer on top of the PECAM1+ endothelial extensions. The cell body of a GFAP+ astrocyte resided directly on the vessel (Fig. 5 I4 and 6), and its extensions were wrapped around the vessel (Fig. 5 G5), albeit not completely ensheathing it (Fig. 5 I2-4).

At E85, claudin-5 increased in intensity and more immunoreactive punctae were present along the vessels, suggesting further strengthening of tight junctions (Fig. 5 J1-J3, K1-K3). The interactions between astrocytes and the vessel wall were even more prominent at E85 (Fig. 5 L1). The astrocyte network around the vessels was denser with only minor gaps (Fig. 5 L2 and L3). In detail, astrocytic processes established close contact to the surrounded vessel (Fig. 5 L4).

### Development of pericyte population

Another essential component of the neurovascular unit are the pericytes marked by the expression of the cell surface tyrosine kinase receptor platelet-derived growth factor receptor β (PDGFRβ) (Smyth et al., 2018). Pericytes are crucial for BBB integrity during embryogenesis, and associate with blood vessels before astrocytes appear. *Pdgfrb^-^/^-^* knockout mice die at birth (Daneman et al., 2010). In adulthood, they cover about one third of the vascular surface (Mathiisen et al., 2010), and the coverage is important for reducing vascular permeability and immune cell entry (Daneman et al., 2010). Being localized within the perivascular space they regulate blood flow velocity and permeability of the capillaries. Recent studies have revealed a surprising link towards enhancing long-term potentiation via activity-evoked pericytic secretion of insulin-like growth factor signaling (Pandey et al., 2023), which is also an important neurotrophin for neural maturation and survival (Benarroch, 2012).

Here, we quantitatively determined the number of PDGFRβ+ cells to gain insight into the distribution and appearance of this cell class (Fig. 6). Cells were scored as positive when they had a clear PDGFRβ+ outline and were closely apposed to a blood vessel (Fig. 6 A, BC). Pericytes frequently resided at the branching point of the vessels suggesting a more efficient sphincter effect at such crossings (Fig. 6 A4 and C4). Already at E60, pericytes were found in all cortical compartments, with a maximum density of 10 cells/mm^2^ in CP/GM and IZ/WM (Fig. 6 D1). At E70, the cell density had increased in all layers, mostly in the CP/GM with 22 cells/mm^2^ (Fig. 6 D2). At E85, an average value of 30 cells/mm^2^ was determined in the CP/GM, while all other layers were still below 25 cells/mm^2^ (Fig. 6 D3). At E100, the cell count continued to increase in all compartments reaching a maximum density of 52 cells/mm^2^ in the CP/GM, with lower densities in the WM and SVZ/VZ (Fig. 6 D4).

**Figure 6.**
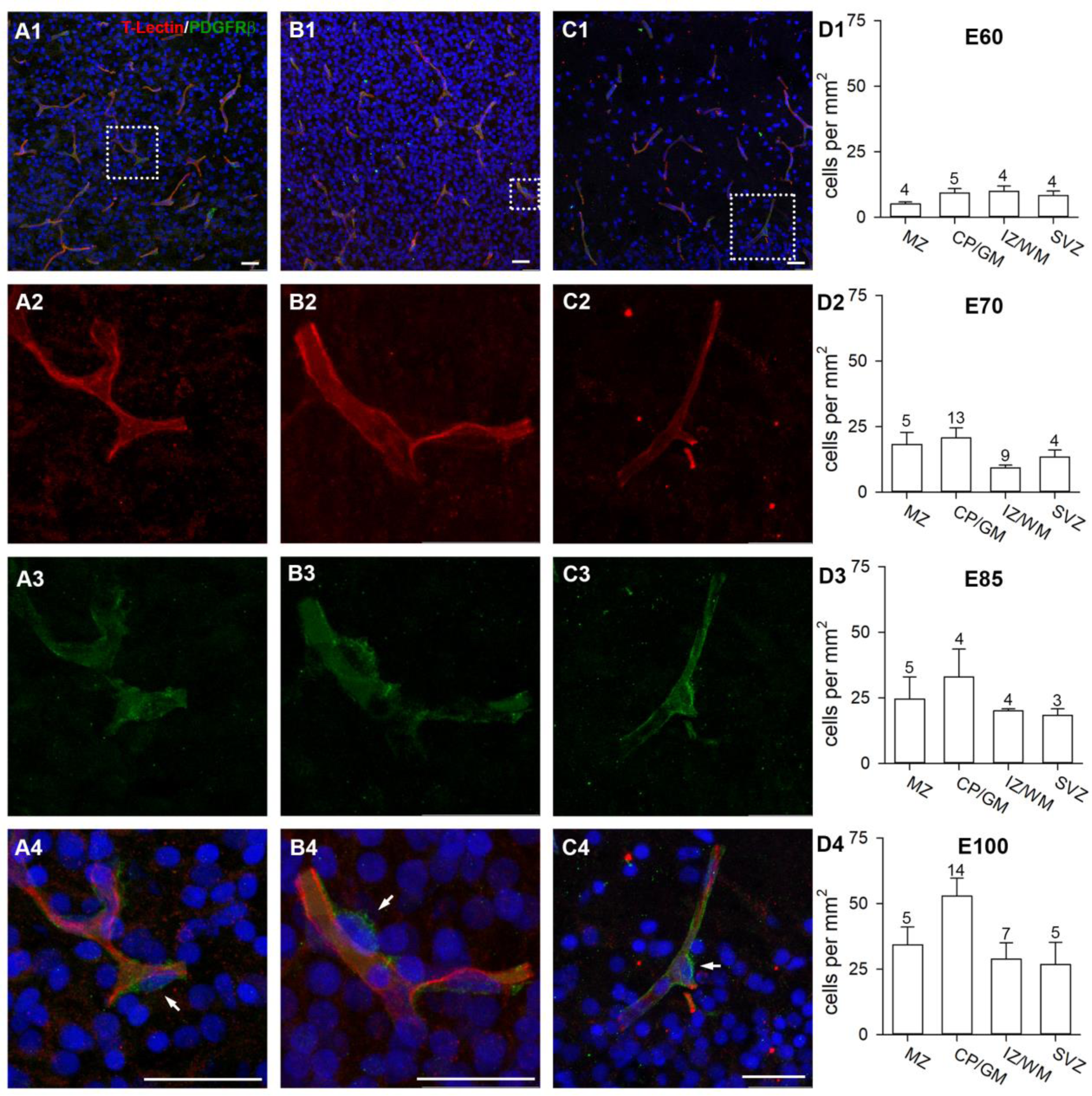
Development of pericytes. Cortex sections were stained with Tomato lectin (red) and PDGFRβ (green). PDGFRβ+ pericytes were plotted in ROI selected by the criterion that every ROI contained at least one pericyte. A1. E60; vessels in L5/6. A2-A4. Magnification of dotted rectangle in A1 showing the single channels and merged. Pericytes (white arrows) were identified as PDGFRβ+ cells forming “bulges” along the vessels. B1. E70; vessels in CP. B2-B4. Magnification of dotted rectangle in B1 showing the single channels and merged. Pericytes have small roundish somata which differ from elongated somata of endothelial cells. C1. E100; vessels in the MZ/L1. C2-C4. Magnification of dotted rectangle in C1 showing the single channels and merged. Pericyte cells often occupy the branching points of blood vessels. D1-D4. Quantification of pericytes from E60 to E100, mean ± S.E.M.. Numbers above the bars report the n of ROIs assessed per laminar compartment. Scale bars: 25 µm.

### Development of vessel-related protein expression in somatosensory and visual cortex

We previously reported that the somatosensory cortex (SC) develops more rapidly than the visual cortex (VC) because the expression of myelin-related proteins occurs earlier (Sobierajski et al., 2023). The finding has been interpreted as being due to the advanced development of the specific somatosensory-evoked activity compared to presumably only spontaneous activity from the retina. This led to the question if the expression of proteins related to angiogenesis also differs between the two sensory areas. Protein expression has been quantified in cortex samples from VC and SC, both containing GM and WM of the apex of gyri with material of E65 being the youngest available in our raw material collection (Fig. 7). Claudin-5 expression was detectable from E65 onwards with increasing intensity (Fig. 7 A), confirming the immunofluorescence stainings (Fig. 5). Expression of AQP4 increased steadily until P90, and the two areas displayed the same expression levels (Fig. 7 B). α-SMA protein was at the adult level in both areas from E65 onwards (Fig. 7 C). The tight junction protein occludin was weakly detectable in 3 out of 5 independent lysates of E65 of the two areas (Fig. 7 D). The major band of predicted 63 kDa was present from E85 onwards, and expression in SC lysates appeared to be subtly accelerated compared to VC. With increasing age, weak side bands of larger molecular weight appeared, possibly representing posttranscriptionally and/or posttranslationally modified forms of occludin (Cummins, 2012). These isoforms were not included in our quantification. The PECAM1/CD31 antibody delivered a single band at 130 kDa (Fig. 7 E). In the younger stages from E60 to E85, the average band intensity was low, it started to increase at E100, and increased further until P90. The cell adhesion molecule vascular endothelial cadherin (VE-cadherin) was well present at E60 followed by a subtle increase until P90 (Fig. 7 F). Finally, comparing only the P90 values obtained for the proteins in VC and SC (corrected for β-actin) did not reveal differences between the two areas. Taken together, the expression of angiogenesis-supporting proteins proceeded almost concurrently in two cortex areas.

**Figure 7.**
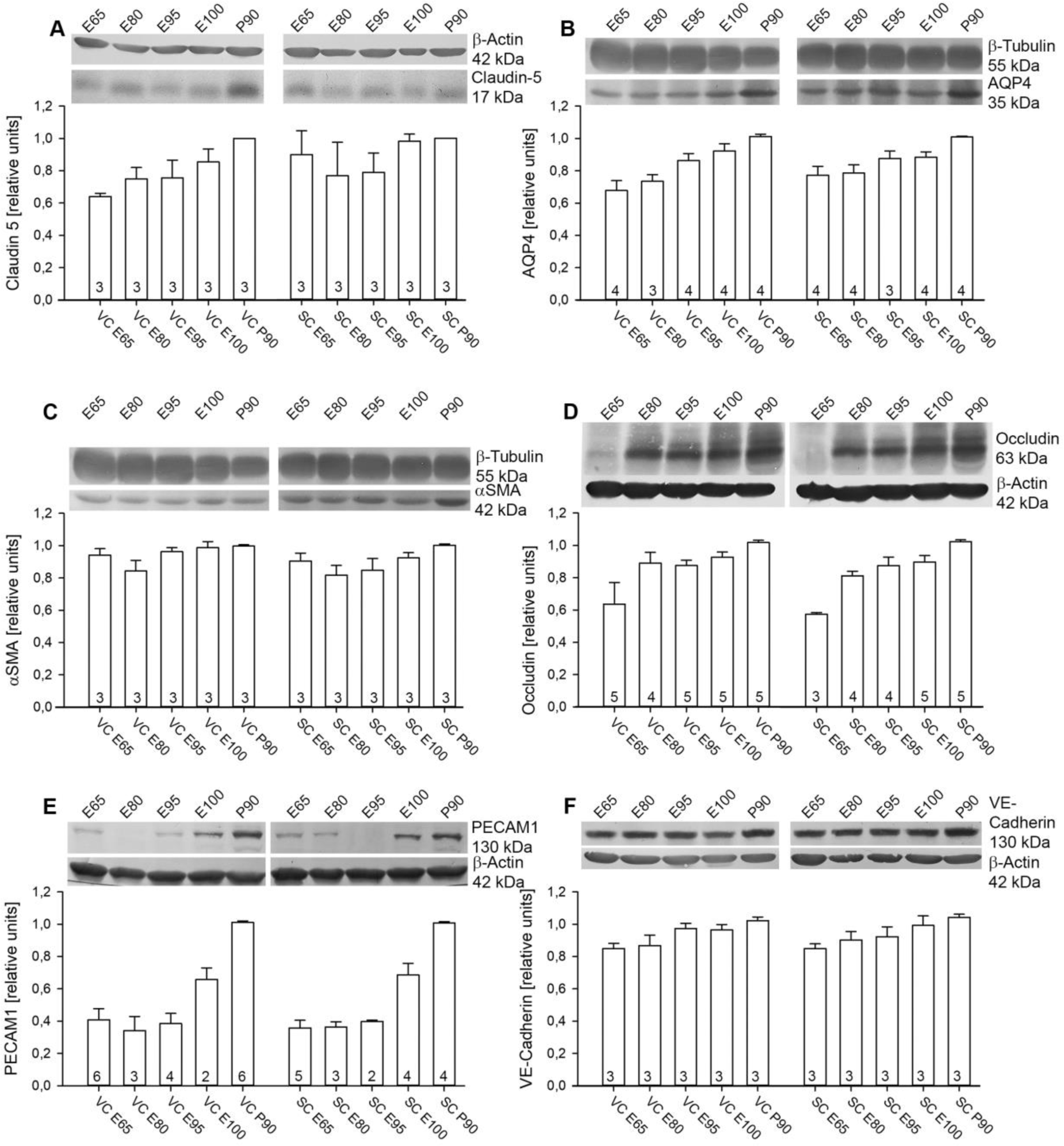
Development of angiogenesis-related proteins. Quantification at E65, E80, E95, E100, and P90 is shown together with representative blots. Visual (VC) and somatosensory (SC) cortex lysates were always run on the same gel to document the area-specific expression profiles. Proteins were normalized to the housekeeping protein β-actin or β-tubulin as indicated. Normalized protein levels were expressed relative to the amount at P90, which was considered adult, and has been set to 1. A. Expression of claudin-5. The 17 kDa band was detected from E65 onwards. B. Expression of AQP4. A 35 kDa band was present at E65 and amounts continuously increased during development. C. Expression of α-SMA at 42 kDa was at near adult levels from E65 onwards. D. Expression of occludin; only the 63 kDa isoform was quantified. Several other bands with larger size appeared with ongoing development. E. Expression of PECAM1 was very low at early ages, gaining intensity at E100. F. Expression of VE-cadherin. A band at the predicted size of 130 kDa was detectable from E65 onwards in lysates of both cortices. The numbers in the bars report the number of independent lysates. Error bars represent mean ± S.E.M.

### Microglia migrate through capillaries and interact with vasculature

As a final component of the neurovascular unit we looked at microglia, the development of which has been quantified previously (Sobierajski et al., 2022). Cells of mesodermal origin colonize the embryonic neuroepithelium as early macrophages, and later differentiate into microglia (Hoeffel and Ginhoux, 2018). This happens before the formation of the BBB is complete. Microglia play important roles e.g. for phagocytosis, synaptic pruning, myelination, and immune surveillance. Recently, it has been shown that microglia cooperate with astrocytes towards phagocytosis of apoptotic cell bodies and neurites (Damisah et al., 2020).

Using ionized calcium-binding adapter molecule 1 (Iba1) as a marker we found round to elongated Iba1+ cell bodies without protrusions within cortical capillaries at E45 in VZ (Fig. 8 A1-4) suggestive of monocyte immigration. Such presumably immigrating cells were observed up to E85 and throughout all laminar compartments (Fig. 8 C1-4). Microglial cells were already in close contact with blood vessels at E68 and extended their processes closely along the capillaries (Fig. 8 B1, B2). During development, the more immature microglial cells are partially IB4+. With maturation towards a fully ramified morphology the IB4 reactivity decreases (Wu et al., 1994). This decline can be recognized in pig between E68 and the E85 stage, the latter containing much more ramified microglia (Sobierajski et al., 2022). However, the punctate IB4+ material associated with Iba1+ microglial cells could also be caused by phagocytosis, since microglia cells contribute to vascular remodeling (Alvarez-Vergara et al., 2021), which is expected to happen more frequently at younger stages. At E85, these close interactions were observed more frequently (Fig. 8 D), and individual processes were wrapped precisely around the vessels (Fig. 8 D1-D3) and rarely contained IB4+ puncta (Fig. 8 D2). This suggested that microglia cells monitor the functionality of the BBB. At P90, the number of microglia observed on and around blood vessels had increased substantially and the cells formed patchy networks around the vessels (Fig. 8 E1-4).

**Figure 8.**
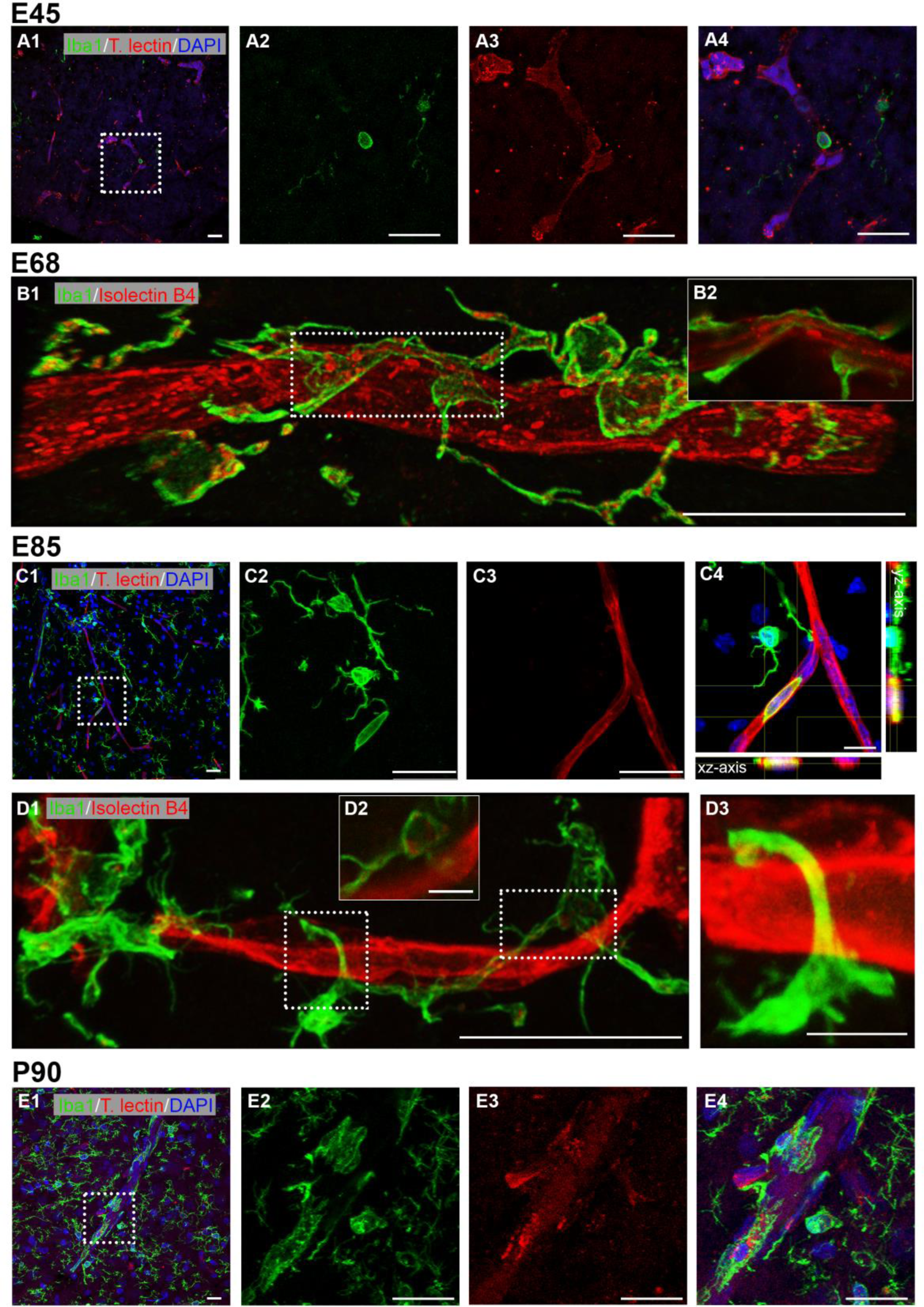
Interaction of microglia and blood vessels. Microglia has been detected with Iba1. Vessels are either stained with tomato lectin or IB4. Microglial cells are also partially IB4+ in particular in more immature stages. A1. Cortex at E45; labeling in VZ. First Iba1+ cells were visible, showing no to few protrusions. A2-A4. Single channels and merged of boxed region in A1 shown at higher magnification. A single microglial cell seemed to emerge from a capillary. B1. Cortex at E68; labeling in GM. Microglial cell wrapping around a vessel. B2. Single z-plane of the boxed region demonstrating that the microglial cell contacts the vessel membrane. Note the frequent association of IB4+ puncta within the Iba1+ microglia which had not yet acquired a fully mature ramified morphology. C1. Cortex at E85; labeling in MZ/L1. C2-C4. Iba1+ monocyte residing inside a blood vessel, as confirmed by xz and yz axis views. D1. Cortex at E85; labeling in L5/6. D2. Single z-plane of the right boxed region in D1, indicating weakly labeled IB4+ puncta associated with the microglial cell. D3. The left boxed region at higher magnification shows a microglial extension wrapping around a vessel. E1. Cortex at P90; microglia-vessel association in the GM. E2-E4. Single channels and merged of boxed region in E1 shown at higher magnification. Scale bars: 25 µm except 10 µm in C4 and 5 µm in D2-D3.

## DISCUSSION

The present study delivers a qualitative and quantitative assessment of the fetal and postnatal development of cerebral cortical vasculature using image analysis of cleared cortex sections and protein blots of two primary sensory areas. At E45 (39% of gestation) a dense network of capillaries could be identified in the midbrain *anlage*, whereas in the cortex primarily layers with high proliferative activity such as the VZ and SVZ already displayed branched capillaries. Penetrating vessels were observed rarely at E45 but more frequently from E70 on. A rapid remodeling then lifted the complexity of the blood vessels network to a near-adult level at E100/110. Additionally, angiogenesis and tight-junction related proteins were expressed at E65 (youngest available) and likely earlier.

### Species comparison of vascular anatomy in cortex

Several previous publications have already delivered quantitative data pools, mostly focusing on rodent models of brain insults or young-to-adult transgenic mice harbouring disease-mediating gene mutations. The pig is of interest as translational model for neurological disorders, most of which are hitherto considered to have neurodevelopmental origins, and many of which manifest early with vascular problems (Ishihara et al., 2023; Ouellette and Lacoste, 2021). As an example, amyloid deposits in microvessels emerge before large plaques appear (Take et al., 2022). Extensive data on fetal and early postnatal development are sparse. Comparisons between studies remained difficult because standard protocols for quantitatively assessing angiogenesis are lacking, and therefore, research teams developed their own procedures up to algorithms and modelling. Further, age, breeding lines, sex, and the cortex area or cortical layer examined varied between studies.

To compare the measures obtained in pig to those published for wildtype healthy rodent, monkey, and human (Table S2, S3). It is also necessary to judge on the variance of data from various sources. Should the variability be enormous and cover the range of the measures obtained in pig, a comparison would not be very meaningful. However, the species comparison revealed rather minor variations (Fig. S2) presumably attributable to natural biological variability, as well as differences in methodologies and analysis strategies. Overall, the blood vessel system occupies approximately the same volume in the species analyzed (Fig. S2 A). Here, the data for mice exhibit the greatest variation, ranging from 0.6 to 3.6% of the volume. The fractional vascular volume ranges between 2-3% in monkey, human, and pig. In neonatal monkey at P2-3 this value is not yet reached, as it is approximately 1%. Moreover, the blood vessel density in relation to the cortical layers is comparable in all species described. There is an overall greater vascular density in GM than in the WM, with highest density in GM layers 5 and 6 (Lauwers et al., 2008; Wu et al., 2014). However, anatomical differences exist (summarized in Fig. S3). The density of vessels in rodents is 1.5-2 times higher compared to density in monkey and human (Fig. S2 B), and pig has a density similar to human. Vessel segment length in rodent is much smaller on average than segment length in pig or human (Table S2). In contrast, the average vessel diameter ranges between 4-5 µm in postnatal rodent and 4-6 µm in E85/P30 pig (Fig. S2 C) whereas vessels of postnatal monkey and human cortex have larger diameters of 7-10 µm.

The greater density and smaller diameter of blood vessels in rodent could be explained by the high relative density of neurons per volume in rodent cortex compared to human cortex. The approximate neuronal density in mice is 0.6 x 10^5^ to 1.6 x 10^5^ neurons/mm^3^, accounting for approximately 40-75% of all cortical cells (Bass et al., 1971; Herculano-Houzel et al., 2006; Irintchev et al., 2005). The neuronal density in human is approximately 0.1 x 10^5^ to 0.5 x 10^5^ neurons/mm^3^, which is substantially lower than in rodent (Gittins and Harrison, 2004; Gredal et al., 2000). A recent EM study reports that the percentage of neurons in human cortex is 40-65% in L2-L4 and 20-30% in L5-L6 (Shapson-Coe et al., 2024). A neuronal density of 0.2 x 10^5^ neurons/mm^3^ is reported for pig visual cortex, which is comparable to that of human (Cragg, 1967). Our data confirm that in P90 pig visual cortex NeuN immunoreactive neurons of all cells are approximately 38% in L2-L4 and 22% in L6 (Sobierajski et al., 2022).

### Assembly of the BBB in cortex

An important question is when the BBB becomes established. A frequently used method has been the injection or perfusion of tracer molecules such as horseradish peroxidase or serum albumin. By injecting human plasma proteins into the vasculature of a 60-day-old sheep fetus (145-150 days of gestation), it has been demonstrated that none of the injected proteins had leaked into the brain, thus, the BBB seemed to be tight and functional at that stage (Dziegielewska et al., 1979). The forebrain of a newborn marsupial, the tammar wallaby (born pea-sized after 27 days), corresponds to the forebrain of a 6-week-old human embryo (Reynolds et al., 1985). Plasma proteins or horseradish peroxidase injected intravenously into newborn tammar wallaby did not penetrate into the brain (Dziegielewska et al., 1988). In the South American opossum, a marsupial born at a very early developmental stage (comparable to an E13 rat) when vessels just begin to form, a <300 kDa tracer does not enter into the brain suggestive of a functional BBB at this very early stage (Ek et al., 2006). In rat, BBB assembly proceeds between E15-17 (Olsson et al., 1968) and in mouse between E13-17 (Risau et al., 1986; Stewart and Hayakawa, 1987). The time course of human BBB assembly relies on detection of the cellular components of the neurovascular unit. During the second trimester in human all components of the unit are present in the pallium including the cerebral cortex (Crouch et al., 2022). In this ensemble of endothelial and mural cell subtypes including smooth muscle cells, pericytes and fibroblasts every cell subtype follows its own developmental trajectory. By the 12^th^ week of gestation, occludin and claudin-5 are found to be expressed in the primary vessels (Virgintino et al., 2000). The appearance of tight junction proteins at this time seems to be sufficient to prevent albumin from entering the brain, providing evidence of early functionality of the BBB. By the 18^th^ week of gestation, occludin and claudin-5 have staining patterns similar to the pattern seen in the adult.

Astrocytes are essential for BBB maintenance. The differentiation of astrocytes and vascular ensheathing begins during the first postnatal week in rodent, and astrocytes appear later than pericytes (Daneman et al., 2010). In human neocortex, first GFAP+ astrocytes are present at the 18^th^ weeks of gestation (Roessmann and Gambetti, 1986). Thus, a certain functionality seems already possible before all components are present, which is probably followed by a “tightening” of the tight junctions and reduction of solute permeability (Hellström et al., 2001). In pig, first evidence of astrocytes ensheating vessels were seen at E45 in midbrain and at E68 in cortex.

Together, the progression of BBB development in pig can be well compared to that in human. At E45, only the tight junction protein claudin-5 could be detected in midbrain and cortex, suggesting an immature state of the BBB. Around mid-pregnancy all cellular and molecular components were present in cortex and the expression levels of the tight junction proteins reached near-adult levels shortly before birth.

### Blood vessel development is not area-specific

Myelin-related proteins show an areal-specific time course of expression (Sobierajski et al., 2023), with advanced expression in the somatosensory cortex where the rostrum is represented. We have suggested that this might have been due to somatosensory-evoked activity. This led to the question of whether the vascular development is also advanced in the somatosensory compared to visual cortex. A study in P1-P60 rat has revealed areal- and laminar-specific differences in vascular density between the visual and auditory cortex and the phylogenetically older entorhinal cortex, but these differences did not emerge before the second postnatal week (Michaloudi et al., 2005). In cat and primate visual cortex, layer 4 in particular develops an increased vascular density postnatally (Fonta and Imbert, 2002; Tieman et al., 2004) reflecting the metabolic requirements of the geniculorecipient layer. We could not see remarkable differences in expression of angiogenesis-related proteins between fetal VC and SC. Recording spontaneous cortical calcium events in prematurely born ferret, before eye and ear opening, has revealed highly similar activity patterns in primary sensory and association areas with functional networks extending over millimeters suggesting that the initial design of the cortical representations follows the same principle (Powell et al., 2024). Such an activity might potentially also promote cortex-wide vascular development which is a prerequisite for the subsequent onset of area-specific neuronal function. Translated to pig, it would support the view that the formation of cortex vasculature during fetal life is independent of the area and the sensory input. Yet remodeling can go on postnatally to adapt the blood supply to the local demand.

## Conflict of interest

The corresponding author, on behalf of the coauthors, declares no conflict of interest.

## Data Availability Statement

All relevant data can be found within the article and its supplementary information.

## Funding Information

This work was supported by Deutsche Forschungsgemeinschaft WA 541/15-1 to PW.

## Acknowledgments

We acknowledge the Regionalverband Ruhr, Essen, Germany, for the interest in our work. We thank Andrea Räk and Christian Riedel for technical support. We thank Prof. Schmelz and Prof. Engelhardt, University Mannheim, Germany, for donating the P5 piglet brain.

## Notes

### Competing Interest Statement

The authors have declared no competing interest.

## REFERENCES

1. Alvarez-Vergara, M. I., Rosales-Nieves, A. E., March-Diaz, R., Rodriguez-Perinan, G., Lara-Ureña, N., Ortega-de San Luis, C., Sanchez-Garcia, M. A., Martin-Bornez, M., Gómez-Gálvez, P. and Vicente-Munuera, P., et al. (2021). Non-productive angiogenesis disassembles Aß plaque-associated blood vessels. Nat Commun 12, 3098. doi:10.1038/s41467-021-23337-z.

2. Andreone, B. J., Lacoste, B. and Gu, C. (2015). Neuronal and vascular interactions. Annu Rev Neurosci 38, 25–46. doi:10.1146/annurev-neuro-071714-033835.

3. Armulik, A., Genové, G., Mäe, M., Nisancioglu, M. H., Wallgard, E., Niaudet, C., He, L., Norlin, J., Lindblom, P. and Strittmatter, K. et al. (2010). Pericytes regulate the blood-brain barrier. Nature 468, 557–561. doi:10.1038/nature09522.

4. Attwell, D., Mishra, A., Hall, C. N., O’Farrell, F. M. and Dalkara, T. (2016). What is a pericyte? J Cereb Blood Flow Metab 36, 451–455. doi:10.1177/0271678X15610340.

5. Bandopadhyay, R., Orte, C., Lawrenson, J. G., Reid, A. R., De, S. S. and Allt, G. (2001). Contractile proteins in pericytes at the blood-brain and blood-retinal barriers. J Neurocytol 30. doi:10.1023/a:1011965307612.

6. Bass, N. H., Hess, H. H., Pope, A. and Thalheimer, C. (1971). Quantitative cytoarchitectonic distribution of neurons, glia, and DNA in rat cerebral cortex. J Comp Neurol 143, 481–490. doi:10.1002/cne.901430405.

7. Battistella, R., Kritsilis, M., Matuskova, H., Haswell, D., Cheng, A. X., Meissner, A., Nedergaard, M. and Lundgaard, I. (2021). Not All Lectins Are Equally Suitable for Labeling Rodent Vasculature. Int J Mol Sci 22. doi:10.3390/ijms222111554.

8. Benarroch, E. E. (2012). Insulin-like growth factors in the brain and their potential clinical implications. Neurology 79, 2148–2153. doi:10.1212/WNL.0b013e3182752eef.

9. Biswas, S., Cottarelli, A. and Agalliu, D. (2020). Neuronal and glial regulation of CNS angiogenesis and barriergenesis. Development 147. doi:10.1242/dev.182279.

10. Böndel, J. C. (2017). Vergleichende morphometrische Untersuchungen am Gehirn von Sus scrofa und Sus scrofa f. domestica, Ludwig-Maximilians-Universität München, Germany.

11. Boscia, F., Esposito, C. L., Casamassa, A., Franciscis, V. de, Annunziato, L. and Cerchia, L. (2013). The isolectin IB4 binds RET receptor tyrosine kinase in microglia. J Neurochem 126, 428–436. doi:10.1111/jnc.12209.

12. Bryant, A., Li, Z., Jayakumar, R., Serrano-Pozo, A., Woost, B., Hu, M., Woodbury, M. E., Wachter, A., Lin, G. and Kwon, T. et al. (2023). Endothelial Cells Are Heterogeneous in Different Brain Regions and Are Dramatically Altered in Alzheimer’s Disease. J Neurosci 43, 4541–4557. doi:10.1523/JNEUROSCI.0237-23.2023.

13. Cassot, F., Lauwers, F., Fouard, C., Prohaska, S. and Lauwers-Cances, V. (2006). A novel three-dimensional computer-assisted method for a quantitative study of microvascular networks of the human cerebral cortex. Microcirculation 13, 1–18. doi:10.1080/10739680500383407.

14. Cayrol, R., Wosik, K., Berard, J. L., Dodelet-Devillers, A., Ifergan, I., Kebir, H., Haqqani, A. S., Kreymborg, K., Krug, S. and Moumdjian, R. et al. (2008). Activated leukocyte cell adhesion molecule promotes leukocyte trafficking into the central nervous system. Nat Immunol 9, 137–145. doi:10.1038/ni1551.

15. Chand, K. K., Miller, S. M., Cowin, G. J., Mohanty, L., Pienaar, J., Colditz, P. B., Bjorkman, S. T. and Wixey, J. A. (2022). Neurovascular Unit Alterations in the Growth-Restricted Newborn Are Improved Following Ibuprofen Treatment. Mol Neurobiol 59, 1018–1040. doi:10.1007/s12035-021-02654-w.

16. Clark, A. T., Abrahamson, E. E., Harper, M. M. and Ikonomovic, M. D. (2022). Chronic effects of blast injury on the microvasculature in a transgenic mouse model of Alzheimer’s disease related Aβ amyloidosis. Fluids Barriers CNS 19, 5. doi:10.1186/s12987-021-00301-z.

17. Cragg, B. G. (1967). The density of synapses and neurones in the motor and visual areas of the cerebral cortex. J Anat 101, 639–654.

18. Crouch, E. E., Bhaduri, A., Andrews, M. G., Cebrian-Silla, A., Diafos, L. N., Birrueta, J. O., Wedderburn-Pugh, K., Valenzuela, E. J., Bennett, N. K. and Eze, U. C. et al. (2022). Ensembles of endothelial and mural cells promote angiogenesis in prenatal human brain. Cell 185, 3753–3769.e18. doi:10.1016/j.cell.2022.09.004.

19. Cummins, P. M. (2012). Occludin: One Protein, Many Forms. Mol Cell Biol 32, 242–250. doi:10.1128/MCB.06029-11.

20. Damisah, E. C., Hill, R. A., Rai, A., Chen, F., Rothlin, C. V., Ghosh, S. and Grutzendler, J. (2020). Astrocytes and microglia play orchestrated roles and respect phagocytic territories during neuronal corpse removal in vivo. Sci Adv 6, eaba3239. doi:10.1126/sciadv.aba3239.

21. Daneman, R., Zhou, L., Kebede, A. A. and Barres, B. A. (2010). Pericytes are required for blood-brain barrier integrity during embryogenesis. Nature 468, 562–566. doi:10.1038/nature09513.

22. Dziegielewska, K. M., Evans, C. A., Malinowska, D. H., Møllgård, K., Reynolds, J. M., Reynolds, M. L. and Saunders, N. R. (1979). Studies of the development of brain barrier systems to lipid insoluble molecules in fetal sheep. J Physiol 292, 207–231.

23. Dziegielewska, K. M., Hinds, L. A., Møllgård, K., Reynolds, M. L. and Saunders, N. R. (1988). Blood-brain, blood-cerebrospinal fluid and cerebrospinal fluid-brain barriers in a marsupial (Macropus eugenii) during development. J Physiol 403, 367–388. doi:10.1113/jphysiol.1988.sp017254.

24. Ek, C. J., Dziegielewska, K. M., Stolp, H. and Saunders, N. R. (2006). Functional effectiveness of the blood-brain barrier to small water-soluble molecules in developing and adult opossum (Monodelphis domestica). J Comp Neurol 496, 13–26. doi:10.1002/cne.20885.

25. Engelhardt, M., Hamad, M. I.K., Jack, A., Ahmed, K., König, J., Rennau, L. M., Jamann, N., Räk, A., Schönfelder, S. and Riedel, C. et al. (2018). Interneuron synaptopathy in developing rat cortex induced by the pro-inflammatory cytokine LIF. Exp Neurol 302, 169–180. doi:10.1016/j.expneurol.2017.12.011.

26. Ernst, L., Darschnik, S., Roos, J., González-Gómez, M., Beemelmans, C., Beemelmans, C., Engelhardt, M., Meyer, G. and Wahle, P. (2018). Fast prenatal development of the NPY neuron system in the neocortex of the European wild boar, Sus scrofa. Brain Struct Funct 223, 3855–3873. doi:10.1007/s00429-018-1725-y.

27. Feeney, J. F. and Watterson, R. L. (1946). The development of the vascular pattern within the walls of the central nervous system of the chick embryo. J Morphol 78, 231–303. doi:10.1002/jmor.1050780205.

28. Fonta, C. and Imbert, M. (2002). Vascularization in the primate visual cortex during development. Cereb Cortex 12, 199–211. doi:10.1093/cercor/12.2.199.

29. Gama Sosa, M. A., Gasperi, R. de, Perez, G. M., Hof, P. R. and Elder, G. A. (2021). Hemovasculogenic origin of blood vessels in the developing mouse brain. J Comp Neurol 529, 340–366. doi:10.1002/cne.24951.

30. Gittins, R. and Harrison, P. J. (2004). Neuronal density, size and shape in the human anterior cingulate cortex: a comparison of Nissl and NeuN staining. Brain Res Bull 63, 155–160. doi:10.1016/j.brainresbull.2004.02.005.

31. Gredal, O., Pakkenberg, H., Karlsborg, M. and Pakkenberg, B. (2000). Unchanged total number of neurons in motor cortex and neocortex in amyotrophic lateral sclerosis: a stereological study. J Neurosci Methods 95, 171–176. doi:10.1016/S0165-0270(99)00175-2.

32. Greene, C., Kealy, J., Humphries, M. M., Gong, Y., Hou, J., Hudson, N., Cassidy, L. M., Martiniano, R., Shashi, V. and Hooper, S. R. et al. (2017). Dose-dependent expression of claudin-5 is a modifying factor in schizophrenia. Mol Psychiatry 23, 2156–2166. doi:10.1038/mp.2017.156.

33. Grubb, S., Cai, C., Hald, B. O., Khennouf, L., Murmu, R. P., Jensen, A. G. K., Fordsmann, J., Zambach, S. and Lauritzen, M. (2020). Precapillary sphincters maintain perfusion in the cerebral cortex. Nat Commun 11, 395. doi:10.1038/s41467-020-14330-z.

34. Hall, C. N., Reynell, C., Gesslein, B., Hamilton, N. B., Mishra, A., Sutherland, B. A., O’Farrell, F. M., Buchan, A. M., Lauritzen, M. and Attwell, D. (2014). Capillary pericytes regulate cerebral blood flow in health and disease. Nature 508, 55–60. doi:10.1038/nature13165.

35. Haruwaka, K., Ikegami, A., Tachibana, Y., Ohno, N., Konishi, H., Hashimoto, A., Matsumoto, M., Kato, D., Ono, R. and Kiyama, H. et al. (2019). Dual microglia effects on blood brain barrier permeability induced by systemic inflammation. Nat Commun 10, 5816. doi:10.1038/s41467-019-13812-z.

36. Haug, H. (1971). Die Membrana limitans gliae superficialis der Sehrinde der Katze. Z Zellforsch 115, 79–87. doi:10.1007/BF00330216.

37. Hellström, M., Gerhardt, H., Kalén, M., Li, X., Eriksson, U., Wolburg, H. and Betsholtz, C. (2001). Lack of pericytes leads to endothelial hyperplasia and abnormal vascular morphogenesis. J Cell Biol 153, 543–553. doi:10.1083/jcb.153.3.543.

38. Henry, V. G. (1968). Fetal Development in European Wild Hogs. The Journal of Wildlife Management 32, 966. doi:10.2307/3799577.

39. Herculano-Houzel, S., Mota, B. and Lent, R. (2006). Cellular scaling rules for rodent brains. Proc Natl Acad Sci U S A 103, 12138–12143. doi:10.1073/pnas.0604911103.

40. Hoeffel, G. and Ginhoux, F. (2018). Fetal monocytes and the origins of tissue-resident macrophages. Cell Immunol 330, 5–15. doi:10.1016/j.cellimm.2018.01.001.

41. Hsu, C.-W., Cerda, J., Kirk, J. M., Turner, W. D., Rasmussen, T. L., Flores Suarez, C. P., Dickinson, M. E. and Wythe, J. D. (2022). EZ Clear for simple, rapid, and robust mouse whole organ clearing. Elife 11. doi:10.7554/eLife.77419.

42. Irintchev, A., Rollenhagen, A., Troncoso, E., Kiss, J. Z. and Schachner, M. (2005). Structural and functional aberrations in the cerebral cortex of tenascin-C deficient mice. Cereb Cortex 15, 950–962. doi:10.1093/cercor/bhh195.

43. Ishihara, K., Takata, K. and Mizutani, K.-I. (2023). Involvement of an Aberrant Vascular System in Neurodevelopmental, Neuropsychiatric, and Neuro-Degenerative Diseases. Life 13. doi:10.3390/life13010221.

44. Ji, X., Ferreira, T., Friedman, B., Liu, R., Liechty, H., Bas, E., Chandrashekar, J. and Kleinfeld, D. (2021). Brain microvasculature has a common topology with local differences in geometry that match metabolic load. Neuron 109, 1168–1187.e13. doi:10.1016/j.neuron.2021.02.006.

45. Kaushik, D. K., Bhattacharya, A., Lozinski, B. M. and Wee Yong, V. (2021). Pericytes as mediators of infiltration of macrophages in multiple sclerosis. J Neuroinflammation 18, 301. doi:10.1186/s12974-021-02358-x.

46. Kaushik, G., Gupta, K., Harms, V., Torr, E., Evans, J., Johnson, H. J., Soref, C., Acevedo-Acevedo, S., Antosiewicz-Bourget, J. and Mamott, D. et al. (2020). Engineered Perineural Vascular Plexus for Modeling Developmental Toxicity. Adv Healthc Mater 9, e2000825. doi:10.1002/adhm.202000825.

47. Kisler, K., Nelson, A. R., Montagne, A. and Zlokovic, B. V. (2017a). Cerebral blood flow regulation and neurovascular dysfunction in Alzheimer disease. Nat Rev Neurosci 18, 419–434. doi:10.1038/nrn.2017.48.

48. Kisler, K., Nelson, A. R., Rege, S. V., Ramanathan, A., Wang, Y., Ahuja, A., Lazic, D., Tsai, P. S., Zhao, Z. and Zhou, Y. et al. (2017b). Pericyte degeneration leads to neurovascular uncoupling and limits oxygen supply to brain. Nat Neurosci 20, 406–416. doi:10.1038/nn.4489.

49. Komabayashi-Suzuki, M., Yamanishi, E., Watanabe, C., Okamura, M., Tabata, H., Iwai, R., Ajioka, I., Matsushita, J., Kidoya, H. and Takakura, N. et al. (2019). Spatiotemporally Dependent Vascularization Is Differently Utilized among Neural Progenitor Subtypes during Neocortical Development. Cell Rep 29, 1113–1129.e5. doi:10.1016/j.celrep.2019.09.048.

50. Kozma, M., Mészáros, Á., Nyúl-Tóth, Á., Molnár, K., Costea, L., Hernádi, Z., Fazakas, C., Farkas, A. E., Wilhelm, I. and Krizbai, I. A. (2021). Cerebral Pericytes and Endothelial Cells Communicate through Inflammasome-Dependent Signals. Int J Mol Sci 22. doi:10.3390/ijms22116122.

51. Kurz, H., Gärtner, T., Eggli, P. S. and Christ, B. (1996). First blood vessels in the avian neural tube are formed by a combination of dorsal angioblast immigration and ventral sprouting of endothelial cells. Dev Biol 173, 133–147. doi:10.1006/dbio.1996.0012.

52. Lauwers, F., Cassot, F., Lauwers-Cances, V., Puwanarajah, P. and Duvernoy, H. (2008). Morphometry of the human cerebral cortex microcirculation: general characteristics and space-related profiles. Neuroimage 39, 936–948. doi:10.1016/j.neuroimage.2007.09.024.

53. Linné, C. von (1758). Systema naturae per regna tria naturae: secundum classes, ordines, genera, species, cum characteribus, differentiis, synonymis, locis. Stockholm, Sweden: Impensis Direct. Laurentii Salvii.

54. Liu, L.-R., Liu, J.-C., Bao, J.-S., Bai, Q.-Q. and Wang, G.-Q. (2020). Interaction of Microglia and Astrocytes in the Neurovascular Unit. Front Immunol 11, 1024. doi:10.3389/fimmu.2020.01024.

55. Mäe, M. A., He, L., Nordling, S., Vazquez-Liebanas, E., Nahar, K., Jung, B., Li, X., Tan, B. C., Chin Foo, J. and Cazenave-Gassiot, A. et al. (2021). Single-Cell Analysis of Blood-Brain Barrier Response to Pericyte Loss. Circ Res 128, e46–e62. doi:10.1161/CIRCRESAHA.120.317473.

56. Mathiisen, T. M., Lehre, K. P., Danbolt, N. C. and Ottersen, O. P. (2010). The perivascular astroglial sheath provides a complete covering of the brain microvessels: an electron microscopic 3D reconstruction. Glia 58, 1094–1103. doi:10.1002/glia.20990.

57. Meyer, G. (2010). Building a human cortex: the evolutionary differentiation of Cajal-Retzius cells and the cortical hem. J Anat 217, 334–343. doi:10.1111/j.1469-7580.2010.01266.x.

58. Michaloudi, H., Grivas, I., Batzios, C., Chiotelli, M. and Papadopoulos, G. C. (2005). Areal and laminar variations in the vascularity of the visual, auditory, and entorhinal cortices of the developing rat brain. Dev Brain Res 155, 60–70. doi:10.1016/j.devbrainres.2004.11.007.

59. Møllgård, K., Beinlich, F. R. M., Kusk, P., Miyakoshi, L. M., Delle, C., Plá, V., Hauglund, N. L., Esmail, T., Rasmussen, M. K. and Gomolka, R. S. et al. (2023). A mesothelium divides the subarachnoid space into functional compartments. Science 379, 84–88. doi:10.1126/science.adc8810.

60. Olsson, Y., Klatzo, I., Sourander, P. and Steinwall, O. (1968). Blood-brain barrier to albumin in embryonic new born and adult rats. Acta Neuropathol 10, 117–122. doi:10.1007/BF00691305.

61. Ouellette, J. and Lacoste, B. (2021). From Neurodevelopmental to Neurodegenerative Disorders: The Vascular Continuum. Front Aging Neurosci 13. doi:10.3389/fnagi.2021.749026.

62. Pandey, K., Bessières, B., Sheng, S. L., Taranda, J., Osten, P., Sandovici, I., Constancia, M. and Alberini, C. M. (2023). Neuronal activity drives IGF2 expression from pericytes to form long-term memory. Neuron. doi:10.1016/j.neuron.2023.08.030.

63. Penna, E., Mangum, J. M., Shepherd, H., Martínez-Cerdeño, V. and Noctor, S. C. (2021). Development of the Neuro-Immune-Vascular Plexus in the Ventricular Zone of the Prenatal Rat Neocortex. Cereb Cortex 31, 2139–2155. doi:10.1093/cercor/bhaa351.

64. Powell, N. J., Hein, B., Kong, D., Elpelt, J., Mulholland, H. N., Kaschube, M. and Smith, G. B. (2024). Common modular architecture across diverse cortical areas in early development. Proc Natl Acad Sci U S A 121, e2313743121. doi:10.1073/pnas.2313743121.

65. Reynolds, M. L., Cavanagh, M. E., Dziegielewska, K. M., Hinds, L. A., Saunders, N. R. and Tyndale-Biscoe, C. H. (1985). Postnatal development of the telencephalon of the tammar wallaby (Macropus eugenii). An accessible model of neocortical differentiation. Anat Embryol 173, 81–94. doi:10.1007/BF00707306.

66. Risau, W., Hallmann, R. and Albrecht, U. (1986). Differentiation-dependent expression of proteins in brain endothelium during development of the blood-brain barrier. Dev Biol 117, 537–545. doi:10.1016/0012-1606(86)90321-0.

67. Ritter, C., Eigen, L., Deiringer, N., Laubscher, L. and Brecht, M. (2023). Coevolution of rostrum and brain in pig species. J Comp Neurol 531, 775–789. doi:10.1002/cne.25461.

68. Roessmann, U. and Gambetti, P. (1986). Astrocytes in the developing human brain. An immunohistochemical study. Acta Neuropathol 70, 308–313. doi:10.1007/BF00686089.

69. Segarra, M., Aburto, M. R., Hefendehl, J. and Acker-Palmer, A. (2019). Neurovascular Interactions in the Nervous System. Annu Rev Cell Dev Biol 35, 615–635. doi:10.1146/annurev-cellbio-100818-125142.

70. Shapson-Coe, A., Januszewski, M., Berger, D. R., Pope, A., Wu, Y., Blakely, T., Schalek, R. L., Li, P. H., Wang, S. and Maitin-Shepard, J. et al. (2024). A petavoxel fragment of human cerebral cortex reconstructed at nanoscale resolution. Science 384, eadk4858. doi:10.1126/science.adk4858.

71. Smyth, L. C. D., Di Xu, Okar, S. V., Dykstra, T., Rustenhoven, J., Papadopoulos, Z., Bhasiin, K., Kim, M. W., Drieu, A.and Mamuladze, T., et al. (2024). Identification of direct connections between the dura and the brain. Nature 627. doi:10.1038/s41586-023-06993-7.

72. Smyth, L. C. D., Rustenhoven, J., Scotter, E. L., Schweder, P., Faull, R. L. M., Park, T. I. H. and Dragunow, M. (2018). Markers for human brain pericytes and smooth muscle cells. J Chem Neuroanat 92, 48–60. doi:10.1016/j.jchemneu.2018.06.001.

73. Sobierajski, E., Lauer, G., Aktas, M., Beemelmans, C., Beemelmans, C., Meyer, G. and Wahle, P. (2022). Development of microglia in fetal and postnatal neocortex of the pig, the European wild boar (Sus scrofa). J Comp Neurol 530, 1341–1362. doi:10.1002/cne.25280.

74. Sobierajski, E., Lauer, G., Czubay, K., Grabietz, H., Beemelmans, C., Beemelmans, C., Meyer, G. and Wahle, P. (2023). Development of myelin in fetal and postnatal neocortex of the pig, the European wild boar Sus scrofa. Brain Struct Funct 228, 947–966. doi:10.1007/s00429-023-02633-y.

75. Stewart, P. A. and Hayakawa, E. M. (1987). Interendothelial junctional changes underlie the developmental ‘tightening’ of the blood-brain barrier. Brain Res 429, 271–281. doi:10.1016/0165-3806(87)90107-6.

76. Stewart, P. A. and Wiley, M. J. (1981). Developing nervous tissue induces formation of blood-brain barrier characteristics in invading endothelial cells: a study using quail--chick transplantation chimeras. Dev Biol 84, 183–192. doi:10.1016/0012-1606(81)90382-1.

77. Take, Y., Chikai, Y., Shimamori, K., Kuragano, M., Kurita, H. and Tokuraku, K. (2022). Amyloid β aggregation induces human brain microvascular endothelial cell death with abnormal actin organization. Biochem Biophys Rep 29, 101189. doi:10.1016/j.bbrep.2021.101189.

78. Tan, X., Liu, W. A., Zhang, X.-J., Shi, W., Ren, S.-Q., Li, Z., Brown, K. N. and Shi, S.-H. (2016). Vascular Influence on Ventral Telencephalic Progenitors and Neocortical Interneuron Production. Dev Cell 36, 624–638. doi:10.1016/j.devcel.2016.02.023.

79. Teng, J., Gao, Y., Yin, H., Bai, Z., Liu, S., Zeng, H., Bai, L., Cai, Z., Zhao, B. and Li, X. et al. (2024). A compendium of genetic regulatory effects across pig tissues. Nat Genet 56, 112–123. doi:10.1038/s41588-023-01585-7.

80. Thomas, J.-L. (2018). Orchestrating cortical brain development. Science 361, 754– 755. doi:10.1126/science.aau7155.

81. Thurgur, H. and Pinteaux, E. (2019). Microglia in the Neurovascular Unit: Blood-Brain Barrier-microglia Interactions After Central Nervous System Disorders. Neuroscience 405, 55–67. doi:10.1016/j.neuroscience.2018.06.046.

82. Tieman, S. B., Möllers, S., Tieman, D. G. and White, J. (2004). The blood supply of the cat’s visual cortex and its postnatal development. Brain Res 998, 100–112. doi:10.1016/j.brainres.2003.11.023.

83. Virgintino, D., Robertson, D., Benagiano, V., Errede, M., Bertossi, M., Ambrosi, G. and Roncali, L. (2000). Immunogold cytochemistry of the blood-brain barrier glucose transporter GLUT1 and endogenous albumin in the developing human brain. Dev Brain Res 123, 95–101. doi:10.1016/S0165-3806(00)00086-9.

84. Weber, B., Keller, A. L., Reichold, J. and Logothetis, N. K. (2008). The microvascular system of the striate and extrastriate visual cortex of the macaque. Cereb Cortex 18, 2318–2330. doi:10.1093/cercor/bhm259.

85. Wu, C. H., Wen, C. Y., Shieh, J. Y. and Ling, E. A. (1994). Down-regulation of membrane glycoprotein in amoeboid microglia transforming into ramified microglia in postnatal rat brain. J Neurocytol 23, 258–269. doi:10.1007/BF01275530.

86. Wu, J., He, Y., Yang, Z., Guo, C., Luo, Q., Zhou, W., Chen, S., Li, A., Xiong, B. and Jiang, T. et al. (2014). 3D BrainCV: simultaneous visualization and analysis of cells and capillaries in a whole mouse brain with one-micron voxel resolution. Neuroimage 87, 199–208. doi:10.1016/j.neuroimage.2013.10.036.

87. Wu, J. Y., Cho, S.-J., Descant, K., Li, P. H., Shapson-Coe, A., Januszewski, M., Berger, D. R., Meyer, C., Casingal, C. and Huda, A. et al. (2024). Mapping of neuronal and glial primary cilia contactome and connectome in the human cerebral cortex. Neuron 112, 41–55.e3. doi:10.1016/j.neuron.2023.09.032.

88. Zilles, K., Palomero-Gallagher, N. and Amunts, K. (2013). Development of cortical folding during evolution and ontogeny. Trends Neurosci 36, 275–284. doi:10.1016/j.tins.2013.01.006.

